# Establishment of split-TurboID in *Plasmodium falciparum* facilitates identification of a PfAlba2 protein interaction network during late blood stage development

**DOI:** 10.64898/2026.11.10.765625

**Authors:** Meghan E. Zadow, Ghizal Siddiqui, Balu Balan, Joshua P. Morrow, Tim Gilberger, Aaron R. Jex, Christopher A. MacRaild, Darren J. Creek, Danny W. Wilson

**Affiliations:** Reseach Centre for Infectious Diseases, School of Biological Sciences, Adelaide University, Adelaide 5005, Australia; Institute for Photonics and Advanced Sensing (IPAS), Adelaide University, Adelaide 5005, Australia; Drug Delivery, Disposition and Dynamics, Monash Institute of Pharmaceutical Sciences, Monash University, Parkville, Victoria 3052, Australia; The Walter and Eliza Hall Institute of Medical Research, Parkville, Australia; University of Melbourne, Melbourne, Australia; Centre for Structural Systems Biology, Hamburg, Germany; Bernhard Nocht Institute for Tropical Medicine, Hamburg; University of Hamburg, Hamburg, Germany; Burnet Institute, Melbourne 3004, Victoria, Australia

## Abstract

Malaria parasites of the genus *Plasmodium* undergo complex multi-host lifecycles that require precise regulation of gene expression. Unlike model eukaryotes, *Plasmodium* spp. genomically encodes a limited repertoire of canonical transcriptional regulators, but possess a substantial complement of RNA-binding proteins (RBPs), highlighting the importance of post-transcriptional regulatory mechanisms in regulating parasite gene expression. Among these RBPs are a family of DNA/RNA-binding proteins termed ‘Alba’ (acetylation lowers binding affinity) of which six have so far been identified in these parasites. Here, we investigated the localisation, functional importance and molecular associations of *P. falciparum* Alba protein PfAlba2 using two different endogenously tagged parasite lines. Using a GFP-FKBP tagged line, we demonstrated that PfAlba2 is expressed throughout asexual blood stage development and primarily localised in the cytosol. Conditional mislocalisation experiments indicated that PfAlba2 contributes to normal parasite growth. To overcome challenges in generating a miniTurboID proximity labelling line, we established a custom split-TurboID proximity labelling system for use in *P. falciparum* to characterise the PfAlba2 proxiome during asexual blood stage development. The split-TurboID system facilitated identification of 101 enriched PfAlba2 proximal proteins in schizont stage development, with significant enrichment for proteins involved in translation initiation, including two additional Alba family members, PfAlba1 and PfAlba4. Complementary RNA immunoprecipitation sequencing (RIP-seq) identified 380 PfAlba2-associated transcripts with significant enrichments including cytosolic ribosomes, rRNA-processing factors, PfAlba1, and egress and invasion-related transcripts (PfRALP1, PfRON4, PfRH1, PfRH5). Together, these findings implicate PfAlba2 in RNA metabolism and translational regulation, suggesting an important regulatory function for this protein during parasite blood stage development.

## Introduction

*Plasmodium* spp. malaria parasites remain a major global health burden, with an estimated ∼282 million malaria cases leading to ∼610,000 deaths in 2024 (WHO, 2025). Despite sustained efforts towards eradication, malaria remains endemic in 80 countries and the majority of infections, severe disease, and deaths are caused by the species *Plasmodium falciparum* (WHO, 2025).

Plasmodium parasites have complex life cycles spanning the human host and mosquito vector. Transmission to humans occurs through the bite of an infected Anopheles mosquito, which introduces sporozoites that infect and replicate in hepatocytes before releasing thousands of progeny into the blood stream where they go on to infect red blood cells and cause symptomatic disease. In *P. falciparum*, blood stage development operates on a continuous ∼48-hour egress and invasion cycle where the parasites undergo 4-5 rounds of asynchronous, asexual replication to produce as many as 32 progeny which are then released into circulation to continue the infection cycle (Figure 1a). A subset of parasites will differentiate into male or female sexual forms (gametocytes) for re-transmission back to the mosquito where they will undergo sexual replication in the midgut.

**Figure 1.**
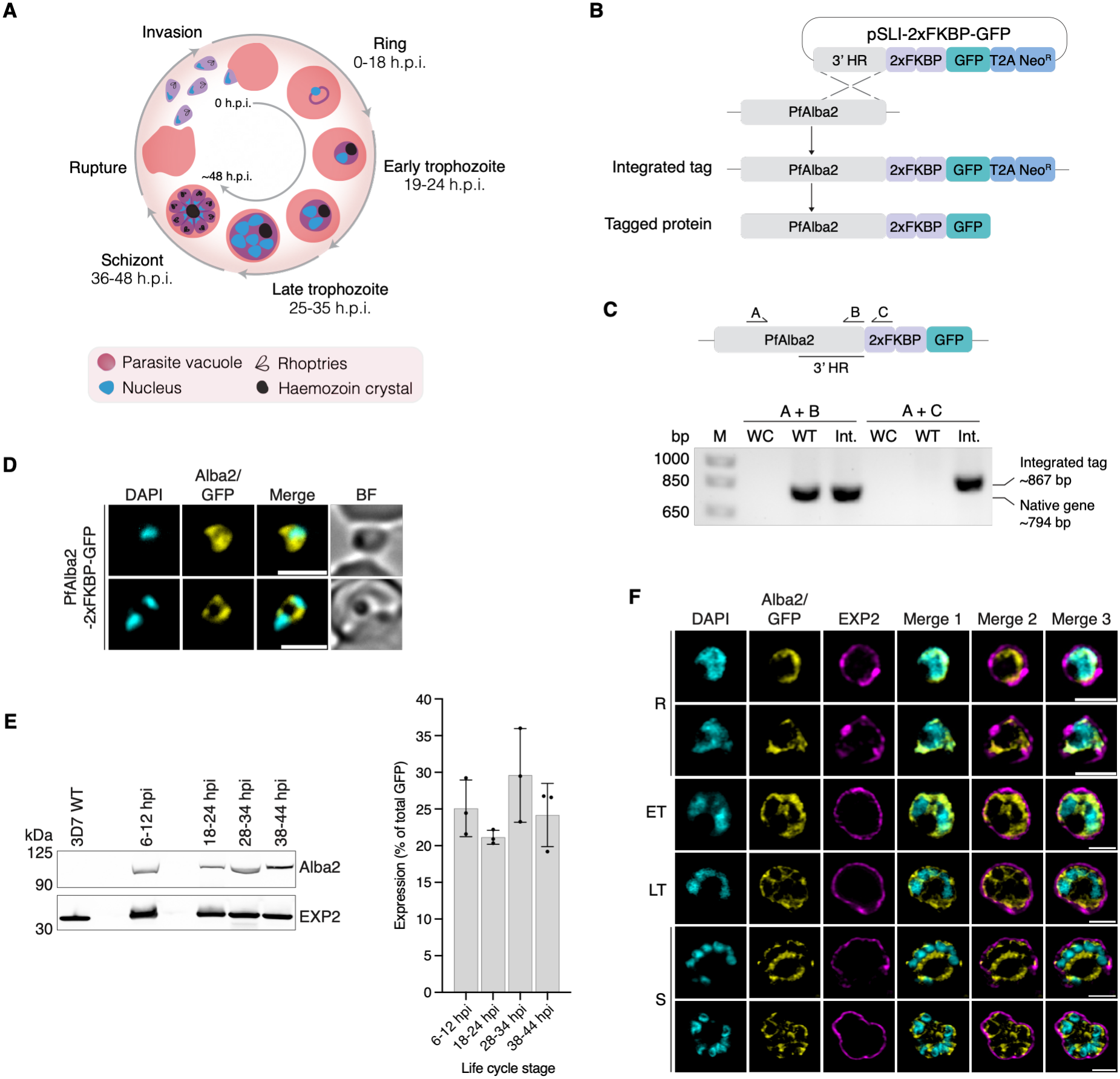
Characterisation of endogenously tagged PfAlba2. **A)** *Plasmodium falciparum* undergoes asynchronous, asexual replication during blood stage development. Parasite growth (ring stage) and replication (trophozoite, schizont stages) occurs over a ∼48-hour period culminating in rupture of the host red blood cell and release of up to ∼32 progeny back into circulation to invade new red blood cells. In this study we utilised a lab-adapted wildtype strain (3D7) which undergoes a slightly shortened ∼44-hour replication cycle. **B)** Endogenous tagging of PfAlba2 was achieved using selection linked integration (SLI) (Birnbaum *et al*, 2017). A C-terminal homology region (HR) for our gene of interest (*Pfalba2*) was inserted into a SLI plasmid vector to facilitate fusion with a 2xFKBP-GFP tag and subsequently transfected into 3D7 wildtype (WT) parasites. Drug treatment was used to first select for episome retention followed by endogenous tag integration. The T2A skip peptide facilitated the expression of a neomycin resistance cassette (Neo^R^) independent from the tagged protein. **C)** Successful endogenous tagging was confirmed by PCR. A region of the native *Pfalba2* gene was amplified by primers A and B (∼794 bp) with primer A corresponding to a region slightly upstream of the 3’HR used in the vector plasmid. Bands of expected size were observed in the 3D7 wildtype (WT) and PfAlba2^2xFKBP-GFP^ (Int.) lanes but not the water control (WC). Primers A and C (where C targets a region of the tag) amplified a ∼867 bp region of the native gene and the endogenous tag which indicated successful integration by presence of a band in the PfAlba2^2xFKBP-GPF^ (Int.) lane only. **D)** Live cell microscopy confirmed expression of the endogenous 2xFKBP-GFP tag. PfAlba2-FKBP-GFP is represented in yellow. Nuclei were stained with DAPI (cyan), merged image represents the GFP and DAPI channels combined, BF=brightfield. Scale bar = 5 μm. **E)** Western blots probed with anti-GFP to detect tagged PfAlba2 (∼90 kDa) showed that PfAlba2 is ubiquitously expressed across blood stage development. 3D7 WT lysate and anti-EXP2 (∼33 kDa) were used as loading controls. PfAlba2 expression at each time point is represented as a percentage of total GFP expression. The dots represent individual data points, columns represent the mean values, and error bars the standard deviation. **F)** Super-resolution microscopy confirmed the largely cytosolic localisation of this protein with some signal also detected at the nuclear periphery, particularly during the early stages of development. Scale bars = 2 μm.

The complex life cycle of Plasmodium parasites are accompanied by highly divergent cell biology compared with many model eukaryotes, with several canonical regulatory pathways and their associated proteins either absent or poorly characterised in *Plasmodium* spp.. This is particularly evident in the regulation of gene expression, where *Plasmodium* spp. possess a relatively limited repertoire of canonical transcriptional regulators. While transcriptional regulation remains important, transcriptomic and proteomic studies have revealed substantial temporal differences between mRNA and protein abundance with ∼30% of transcripts exhibiting delayed protein expression during asexual blood stage development (Caro *et al*, 2014; Foth *et al*, 2011; Foth *et al*, 2008; Le Roch *et al*, 2004). These observations, together with the high abundance of RNA-binding proteins (RBPs) encoded by Plasmodium parasites, suggests a greater reliance on post-transcriptional regulation in coordinating gene expression (Reddy *et al*, 2015).

Among the RBPs implicated in post-transcriptional regulation are members of the acetylation lowers binding affinity (Alba) protein family, which has undergone expansion in numbers in malaria parasites relative to their compact genomes (recently reviewed in Zadow *et al* (2025)). Six Alba domain-containing proteins have so far been identified in *P. falciparum*and are annotated as PfAlba1-6 (Chêne *et al*, 2012; Reddy *et al*., 2015). Alba proteins are conserved across all clades in the phylogenetic tree of life and are characterised by a highly conserved structural domain consisting of beta-sheets and alpha-helices (Guerrier-Takada *et al*, 2002; Li & Altman, 2001). Alba proteins were previously assigned to one of three phylogenetic lineages including two eukaryotic lineages: Rpp20/Pop7 and Rpp25/Mdp2. However, a more recent phylogenetic study has expanded this into 13 distinct clusters, further highlighting the evolutionary diversity of this protein family (Aravind *et al*, 2003; Jagadeesh & Vembar, 2024). Beyond the conserved Alba domain, many of these proteins possess additional domains that may contribute to the broad functional diversity seen within this protein family (Goyal *et al*, 2016). Consistent with this diversity, Alba proteins are reported to be involved in critical cellular processes including the regulation of key virulence genes, response to stress, transitions between hosts, and stage differentiation (reviewed in Zadow *et al*. (2025)).

Despite several studies investigating PfAlba proteins, their functional characterisation has been limited by challenges in generating genetically modified parasite lines. Consequently, previous studies have relied upon in-house antibodies and episomal overexpression systems, limiting opportunities for direct functional analysis of these proteins (Acharya *et al*, 2025; Chêne *et al*., 2012; Vembar *et al*, 2015). These studies have suggested that PfAlba proteins are essential to the parasite blood stage, consistent with a genome wide saturation mutagenesis study that identified several PfAlba genes as potentially essential for parasite blood stage development (Zhang *et al*, 2018).

Release of the first Plasmodium genome in 2002, together with rapid advancements in gene-editing in recent years, has substantially improved our knowledge on the function and importance of individual parasite proteins (Bushell *et al*, 2017; de Koning-Ward *et al*, 2015; Gardner *et al*, 2002; Zhang *et al*., 2018). However, proteins rarely function in isolation, and understanding their interactions with other proteins, and other substrates, such as RNA, is essential to elucidate their role in parasite biology.

Proteomics approaches such as affinity purification and proximity labelling (BioID, TurboID) have become valuable tools for investigating the cell biology and protein interactomes of malaria parasites. Affinity capture techniques, also known as coimmunoprecipitation (CoIP), enable isolation of protein complexes associated with a bait protein, typically through antibody capture (Dunham et al, 2012). In contrast, proximity-labelling approaches such as TurboID and miniTurboID utilise a modified BirA* biotin ligase to biotinylate proteins in specific subcellular niches or proximal to a bait protein upon biotin supplementation (Branon *et al*, 2018a; Roux *et al*, 2012). Proximity labelling methods compatible with *Plasmodium* have previously been reviewed by Kimmel *et al* (2022).

More recently, a new iteration of the BirA* proximity labelling system has been developed termed ‘Split-TurboID’ or ‘ContactID’ (Cho et al, 2020a; Cho et al, 2020b; De Munter et al, 2017; Kwak et al, 2020; Schopp et al, 2017). This alternative TurboID system involves splitting the BirA* enzyme into two inactive fragments that only regain activity upon reconstitution in the presence of a dimeriser. This inducible biotinylation option can offer significantly lower background, increased specificity and lower cell toxicity compared to TurboID and miniTurboID options which feature constitutive BirA* activity. While not yet established in Plasmodium parasites, split-TurboID offers a valuable addition to the proximity labelling toolkit for these unique organisms.

Previous studies have reported difficulties in endogenously tagging *P. falciparum* Alba proteins, limiting their direct functional characterisation (Acharya *et al*., 2025; Chêne *et al*., 2012). However, we recently successfully tagged PfAlba4 and used this line to characterise this Alba protein’s function, including identification of PfAlba4’s proximal proteins (proxiome) using miniTurboID (Review Commons MS RC-2026-03833, referred to as *manuscript submitted* from now on). Building on this work, we generated endogenously tagged PfAlba2 parasite lines to examine its localisation, functional importance, and RNA associations during asexual blood stage development. To overcome challenges associated with miniTurboID proximity labelling for PfAlba2, we additionally established a customised split-TurboID system for use in *P. falciparum* that identified a high confidence proxiome for PfAlba2 and also provides a robust addition to the toolkit of proximity labelling options for use in these highly divergent organisms. Together, our findings reveal associations between PfAlba2, translation-associated proteins, and transcripts involved in RNA metabolism and parasite invasion, supporting a potential role for PfAlba2 in post-transcriptional regulation during asexual blood-stage development.

## Results

### PfAlba2 is a ubiquitously expressed nucleocytoplasmic protein important for blood stage development

PfAlba2 is a 25 kDa protein consisting of an Alba domain followed by an RGG/RG repeat RNA-binding domain and belongs to the Rpp25-like lineage of Alba domain proteins (Aravind *et al*., 2003; Jagadeesh & Vembar, 2024). PfAlba2 is predicted to be essential to the parasite blood stage based on a genome-wide saturation mutagenesis screen and previous unsuccessful attempts to endogenously tag this protein (Zhang *et al*., 2018).

In a recent study, episomal overexpression of PfAlba2 was shown to disrupt mRNA homeostasis and impair the progression of parasite growth from rings to late stage parasites (Acharya *et al*., 2025). PfAlba2 has also been implicated in virulence gene regulation in *P. falciparum* and processes involved in host transitions in the rodent malaria parasites *P. berghei* and *P. yoelii* (Acharya *et al*., 2025; Mair *et al*, 2010; Muñoz *et al*, 2017).

To investigate PfAlba2 function, we first generated a 2xFKBP-GFP endogenously tagged parasite line using selection-linked integration (PfAlba2^2xFKBP-GFP^, Figure 1b) (Birnbaum *et al*, 2017). Successful integration of the tag at the C-terminus of 3D7 PfAlba2 was confirmed by PCR (Figure 1c). Expression of the tagged protein was then confirmed by live cell microscopy and western blot, which showed that PfAlba2 is expressed ubiquitously across blood stage development (Figure 1d, e; Supplementary figure 1a; Supplementary data 1). Using super-resolution microscopy, we observed that PfAlba2 has a largely cytoplasmic localisation throughout the blood stage but also shows some overlap with the parasite nucleus, particularly during the early stages of development (Figure 1f; Supplementary figure 1b). This localisation and expression are consistent with previous studies which utilised over-expression vectors or in-house antibodies to study this protein (Acharya *et al*., 2025; Chêne *et al*., 2012).

We next investigated the impact of PfAlba2 mislocalisation across blood stage development utilising an inducible knock-sideways (KS) approach to sequester the protein in the nucleus at different developmental stages (Figure 2a) (Birnbaum *et al*., 2017). To facilitate this, we co-transfected the PfAlba2^2xFKBP-GFP^ line with an episome expressing the dimerisable FRB subunit linked to a nuclear mislocalisation signal (PfAlba2^2xFKBP-GFP/NLS-FRB-mCh^). Retention and expression of the episome was confirmed by western blot and live cell microscopy (Figure 2b; Supplementary figure 1c). Rapalog-inducible mislocalisation in the PfAlba2-KS line was confirmed by live cell microscopy where an overlapping of GFP (PfAlba2) and mCherry (NLS) signals was observed within the nucleus in the presence of rapalog (Figure 2c; Supplementary figure 1c).

**Figure 2.**
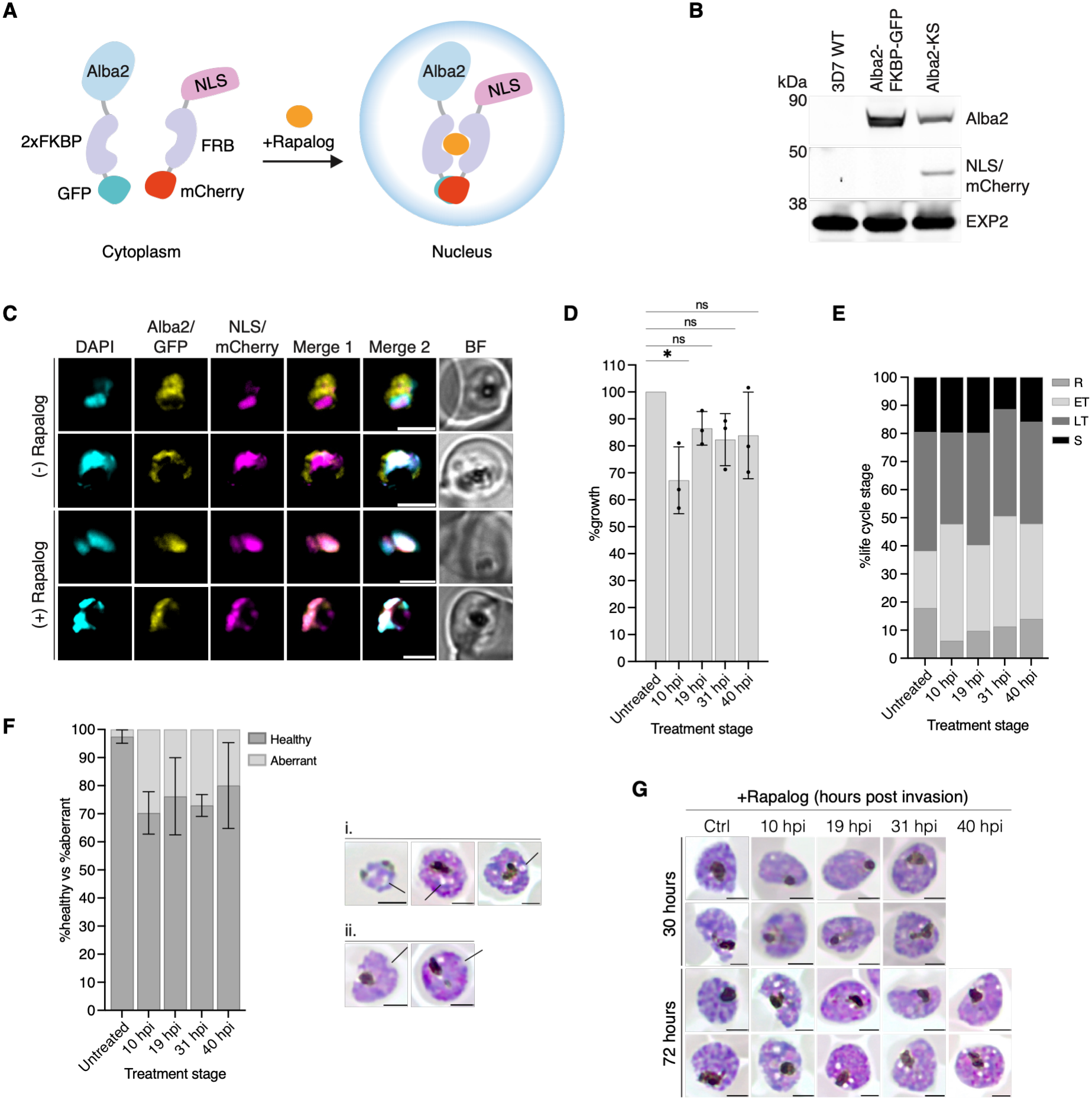
Nuclear mislocalisation of PfAlba2 during blood stage development has a moderate impact on parasite growth. **A)** Schematic representation of the conditional mislocalisation/knock-sideways (KS) strategy. An episome containing a nuclear mislocalisation signal (NLS) fused with a dimerisable FRB subunit and mCherry was co-transfected into the PfAlba2^2xFKBP-GFP^ transgenic line. Addition of rapalog induces dimerisation between the FKBP fused with PfAlba2 and the episomally expressed FRB subunit fused with the NLS to facilitate translocation of PfAlba2 into the nucleus and away from its likely site of function, the cytosol. **B)** Successful co-transfection was confirmed by western blots probed with anti-mCherry to detect the episome. Anti-GFP was used to detect the endogenously tagged PfAlba2 and a band of expected size (∼90 kDa) is present in the PfAlba2-FKBP-GFP control and the PfAlb2-KS sample but absent in the 3D7 WT control lysate. An mCherry band indicates presence of the mislocalisation plasmid (∼45 kDa) and is only observed in the PfAlba2-KS lane. Anit-EXP2 was used as a loading control (∼33 kDa). **C)** Live cell microscopy analysis in the presence (+) or absence (-) of rapalog demonstrates efficient dimerisation and mislocalisation. Nuclei were stained with DAPI (cyan), PfAlba2-FKBP-GFP is represented in yellow, and the episome in magenta (NLS-mCherry). Merge 1 – GFP and mCherry, merge 2 – DAPI, GFP, mCherry. BF = brightfield and scale bars = 5 μm. **D)** Nuclear mislocalisation of PfAlba2 had a moderate impact on parasite growth across blood stage development. Cultures of PfAlba2^2xFKBPGFP/NLS-FRB-mCh^ parasites were treated with rapalog to induce PfAlba2 mislocalisation at specific time points during blood stage development – ring stage (10 h.p.i.), early trophozoites (19 h.p.i.), late trophozoites (31 h.p.i.), and schizonts (40 h.p.i.) (h.p.i. = hours post infection). The rapalog was removed 48-hours post-assay set-up and parasite growth measured at 72-hours. Growth of rapalog-treated parasites is represented as a percentage compared to the untreated control parasites. The mean of n=3 independent replicates is shown. Significance was determined by two-tailed paired t-tests (*\*p* <0.05, ns = not significant) and error bars for all graphs show standard deviation. **E)** Mislocalisation to the nucleus did not significantly impact distribution of life stages present at 72-hours post assay set-up however, **F)** an increase in aberrant phenotypes was observed, predominantly consisting of parasites with stress vacuoles and apparent cellular disorganisation (i) and membrane irregularities (ii). **G)** Giemsa images representative of parasites observed in each treatment group at 30- and 72-hours post assay set-up. Scale bars = 2 μm.

Conditional nuclear mislocalisation of PfAlba2 during the blood stage appeared to impact parasite growth, with significant growth reduction observed in PfAlba2-KS parasites where rapalog-mediated mislocalisation was induced at ring-stages (10 h.p.i.) (Figure 2d; Supplementary data 1). While there were no obvious differences in developmental progression post rapalog treatment, an increase in aberrant parasites was observed for all treatment groups compared to the untreated control at the final timepoint (72 hours), with up to 30% of parasites displaying stress vacuoles, membrane irregularities and/or cellular disorganisation (Figure 2e-g; figure 2a, b). Additional control assays using 3D7 wildtype parasites were also undertaken and these assays indicated no significant impact due to the rapalog treatment alone (Supplementary figure 2c-f; Supplementary data 1).

Overall, these data are congruent with previous reports of PfAlba2 expression and localisation, and knock-sideways assays indicate a potentially important role for this protein during parasite blood stage development. These observations are similar to our findings for another PfAlba protein, PfAlba4, which also exhibited ubiquitous expression and cytoplasmic localisation throughout blood stage development and a similar, though more pronounced, knock-sideways phenotype, particularly in ring stage parasites (Zadow *et al*, *Manuscript submitted*).

### Development of a split-TurboID system for use in *Plasmodium* parasites

Previous work in our lab has utilised an episomally expressed miniTurboID proximity labelling system to investigate the protein interaction network of the related DNA/RNA-binding protein PfAlba4 (Zadow *et al*, *Manuscript submitted*). The miniTurboID episome features a constitutively active BirA* biotin ligase fused to a dimerisable subunit (FRB) and facilitates biotinylation of proteins proximal to a bait protein in the presence of the dimeriser rapalog. However, repeated attempts to generate a similar line for PfAlba2 were unsuccessful, with the parasite unable to maintain the miniTurboID episome during prolonged culture, potentially due to the high activity and indiscriminate biotinylation by the BirA* enzyme. To overcome this limitation, we adopted a rapalog-inducible split-TurboID proximity labelling system (Figure 3a), an approach not previously established in *Plasmodium* parasites to our knowledge (Cho *et al*., 2020a; Kwak *et al*., 2020; Schopp *et al*., 2017).

**Figure 3.**
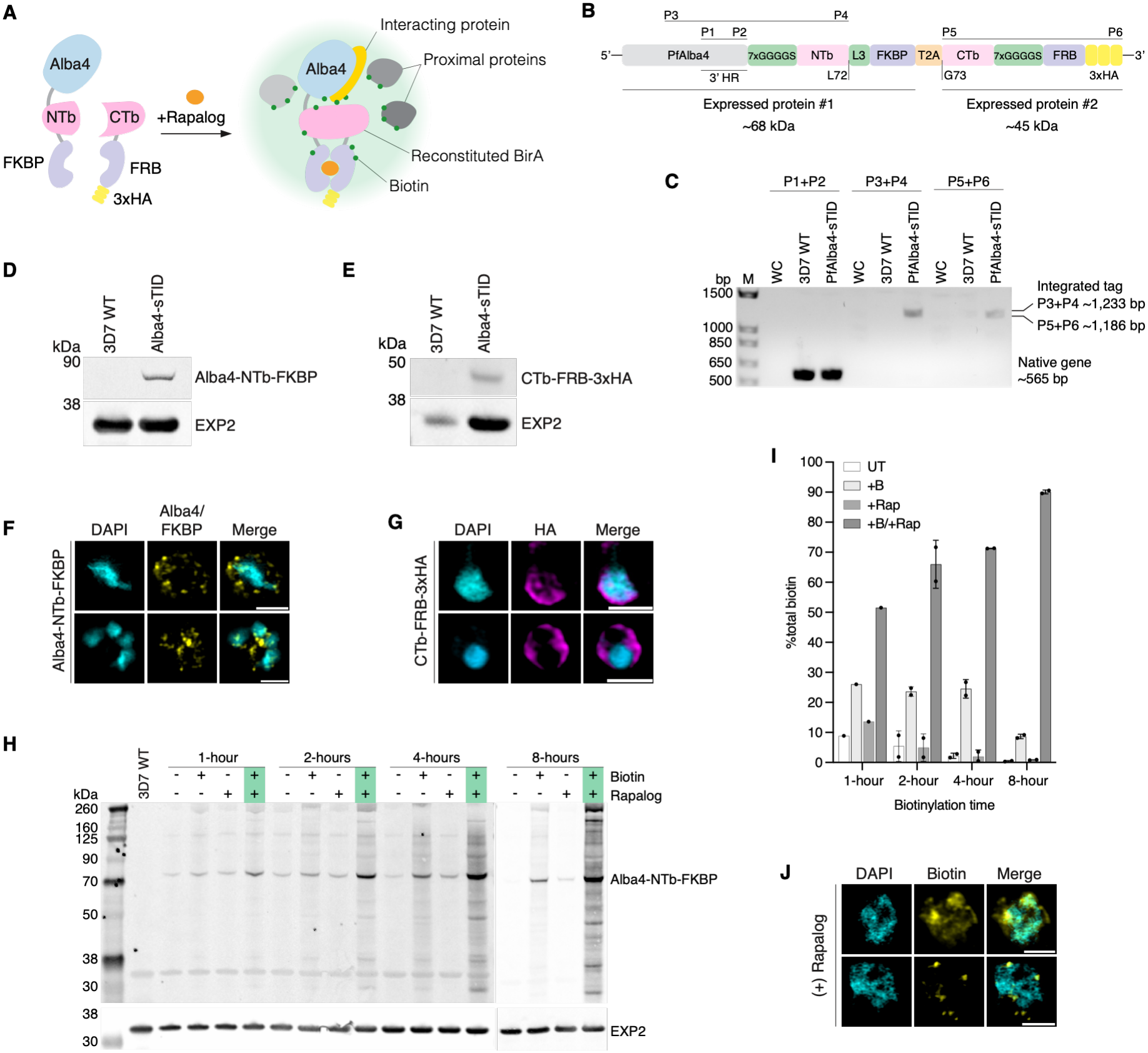
Development and validation of a split-TurboID system for use in *P. falciparum*. **A)** Schematic representation of the split-TurboID proximity labelling strategy. A split-TurboID tag is appended to a target protein (PfAlba4). Upon translation of the tagged target protein, the N-terminal portion of the tag (NTb-FKBP) remains fused with the target protein while the C-terminal portion (CTb-FRB-3xHA) is translated as a separate protein. Reconstitution and activity of the BirA* is induced by the addition of the drug rapalog, facilitating inducible biotinylation of proximal proteins. **B)** Schematic of the split-TurboID endogenous tag. 7xGGGGS and L3 are linking regions. The 3’ homology region (HR) of PfAlba4 used to generate the plasmid is indicated, L72 and G73 refer to the amino acid residues of the TurboID sequence where the protein was separated. Endogenously tagged lines were achieved using selection linked integration as previously outlined (Figure 1b) (Birnbaum *et al*., 2017) and P1-P6 indicate primers used to confirm tag integration. **C)** Successful tag integration was confirmed by PCR. Water controls (WC) and 3D7 WT controls were used for each primer pair, no bands were observed for any of the WC. P1 and P2 correspond to the native *Pfalba4* gene sequence and bands of ∼565 bp were observed for the 3D7 WT and PfAlba4-sTID gDNA as expected. P3 corresponds to a region of the native gene upstream of the 3’HR used to generate the transfection plasmid and P4 corresponds to a sequence in the NTb sequence. For these primers, a ∼1,233 bp band was only observed in the PfAlba4-sTID lane, indicating successful integration of the tag. Primers P5 and P6 amplify the 3’ region of the tag and the expected ∼1,186 bp band is once again only observed in the PfAlba4-sTID lane and absent in controls, indicating this sequence is only present in the tag line. **D)** Western detection of split-TurboID expression. A band corresponding to endogenously tagged PfAlba4-NTb-FKBP was detected at ∼73 kDa by probing with anti-FKBP and, **E)** presence of the CTb-FRB-3xHA fragment was confirmed by probing with anti-HA and showed a corresponding band of expected size at ∼45 kDa. 3D7 WT lysate and anti-EXP2 were used as loading controls. **F)** Immunofluorescence microscopy showed the localisation of the two split-TurboID components. PfAlba4-NTb-FKBP was detecting by an anti-FKPB antibody (Yellow) and **G)** the CTb-FRB-3xHA component was detected with anti-HA (magenta). Both split-Turbo proteins were detected in the cytoplasm and localisation of PfAlba4-NTb-FKBP was consistent with previous localisation for this protein. Nuclei were stained with DAPI (cyan), scale bars = 2 μm. **H)** Representative western blot probed with streptavidin to detect biotinylated proteins in PfAlba4-sTID cultures grown in biotin-free media. Samples were incubated for 1-, 2-, 4-, or 8-hours incubation under the following conditions: untreated control (-/-), +Biotin only control (+/-), +Rapalog only control (-/+) and +Biotin/+Rapalog treated samples (+/+). Bands indicate biotinylated proteins and a band corresponding to PfAlba4-NTb-FKBP was observed at ∼73 kDa. 3D7 WT lysate and anti-EXP2 were used as loading controls. **I)** Quantified western blot streptavidin signal showing level of biotinylation in untreated (UT), +Biotin only (+B), +Rapalog only (+Rap), and +Biotin/+Rapalog (+B/+Rap) samples. Signal was normalised across lanes using the EXP2 control and biotinylation was calculated as a percentage of the total streptavidin signal for each time point. Dots represent the individual data points, columns show the mean, and standard deviation is represented by the error bars. **J)** Biotinylation in the presence (+) of rapalog and exogenous biotin was confirmed by immunofluorescence microscopy. Coverslips were stained with DAPI (nuclei, cyan) and streptavidin (biotin, yellow). The biotin signal appears largely cytosolic, as expected for PfAlba4, and indicates activity of the reconstituted BirA* enzyme. Scale bars = 2 μm.

We designed a split-TurboID tagging system that could be appended to the C-terminus of a target with a single transfection utilising existing selection-linked integration tagging methods (Figure 3b) (Birnbaum *et al*., 2017). The construct incorporated the low-activity BirA* split site, L72/G73, previously identified by Cho *et al*. (2020a). Although the entire tag is integrated at the C-terminus of the target protein, the inclusion of a T2A ribosomal skip peptide facilitates the separate expression of the N-terminal TurboID (NTb) fused to the target protein and FKBP, and the C-terminal fragment (CTb) fused to the FRB subunit and 3x haemagglutinin (HA) tag.

To establish the utility of the split-TurboID tag, and to allow comparison with our existing PfAlba4-miniTurboID system, we first generated a PfAlba4 split-TurboID parasite line (PfAlba4^NTb-FKBP/CTb-FRB-3xHA^) by introducing the construct into existing selection-linked integration plasmids and transfecting into 3D7 wildtype parasites. PCR was used to show successful integration of the tag into the parasite genome (Figure 3c) and western blot detection of the FKBP and HA tag components confirmed PfAlba4-NTb-FKBP tagged protein expression, as well as the independent expression of the CTb-FRB-3xHA tag (Figure 3d, e). Immunofluorescence microscopy showed that the PfAlba4-NTb-FKBP protein localises to the cytoplasm, consistent with expectations for PfAlba4 (Figure 3f; Supplementary figure 3a). Microscopy analysis further indicated that the CTb-FRB-3xHA also remains in the cytoplasm after translation (Figure 3g; Supplementary figure 3b).

To confirm function and determine biotinylation efficiency of the PfAlba4-sTID line we harvested sequential cultures after 1-, 2-, 4-, and 8-hours of incubation under the following conditions – Untreated, biotin only (+Biotin), Rapalog only (+Rapalog), biotin and Rapalog (+B/+R). These experiments were repeated for cultures grown in regular HEPES-RPMI (h-RPMI) culture media which contains endogenous biotin, and cultures grown in biotin-free media. For parasites grown in h-RPMI media, biotinylation was observed across all samples, even in the absence of the dimeriser rapalog. Given that the split-TurboID system relies on rapalog-inducible dimerisation for restoration of function, this may indicate some level of rapalog-independent interaction between the FKBP/FRB subunits and thus, the split-TurboID fragments (Supplementary figure 4a). However, longer biotinylation times did increase the proportion of biotinylation present in the +B/+R samples compared to +Biotin only controls, indicating induced dimerisation of the BirA* increased biotinylation efficiency well above that of the non-dimerised BirA* with longer incubation times (Supplementary figure 4b; Supplementary data 2).

For the PfAlba4-sTID parasites incubated in biotin-free media there was minimal biotinylation present in the untreated and +Rapalog samples at all time points and the band that was most strongly biotinylated was one consistent with the size of PfAlba4-NTb-FKBP (Figure 3h, i; Supplementary figure 5). Quantitative western blots showed that by 4-hours incubation, ∼70% of biotinylation was occurring in the +B/+R treated sample, and by 8-hours this had reached ∼90% (Figure 3i; Supplementary data 2). Biotinylation in the biotin-only control remained below 30% for each time point and had dropped to ∼10% by 8-hours incubation. A band consistent with the expected size of PfAlba4-NTb-FKBP was clearly observable in all biotin-treated samples, particularly in the +B/+R samples, indicating the increased biotinylation in the presence of rapalog-induced dimerisation of the split-TurboID (Figure 3h).

Finally, we performed immunofluorescence assays to visualise biotinylation in the presence of rapalog. A mostly cytoplasmic biotinylation signal was observed in these cells, consistent with the localisation of PfAlba4, further indicating efficient dimerisation and biotinylation with the protein of interest. (Figure 3j; Supplementary figure 3c). Together, these data indicate the efficient, rapalog-inducible biotinylation of this custom split-TurboID system.

### Split-TurboID identifies a high confidence proxiome for PfAlba4 that overlaps with previous PfAlba4-BioID experiments

To further confirm the utility and specificity of the split-TurboID, we undertook proteomics experiments using the PfAlba4-sTID parasite line for comparison against a previous PfAlba4 proximity labelling experiment which utilised an episomally expressed miniTurboID (mTID) system (Zadow *et al*, *Manuscript submitted*). Biotinylation quality for all replicates was validated by western blot prior to subsequent downstream processing (Figure 4a; Supplementary figure 6a, b)

**Figure 4.**
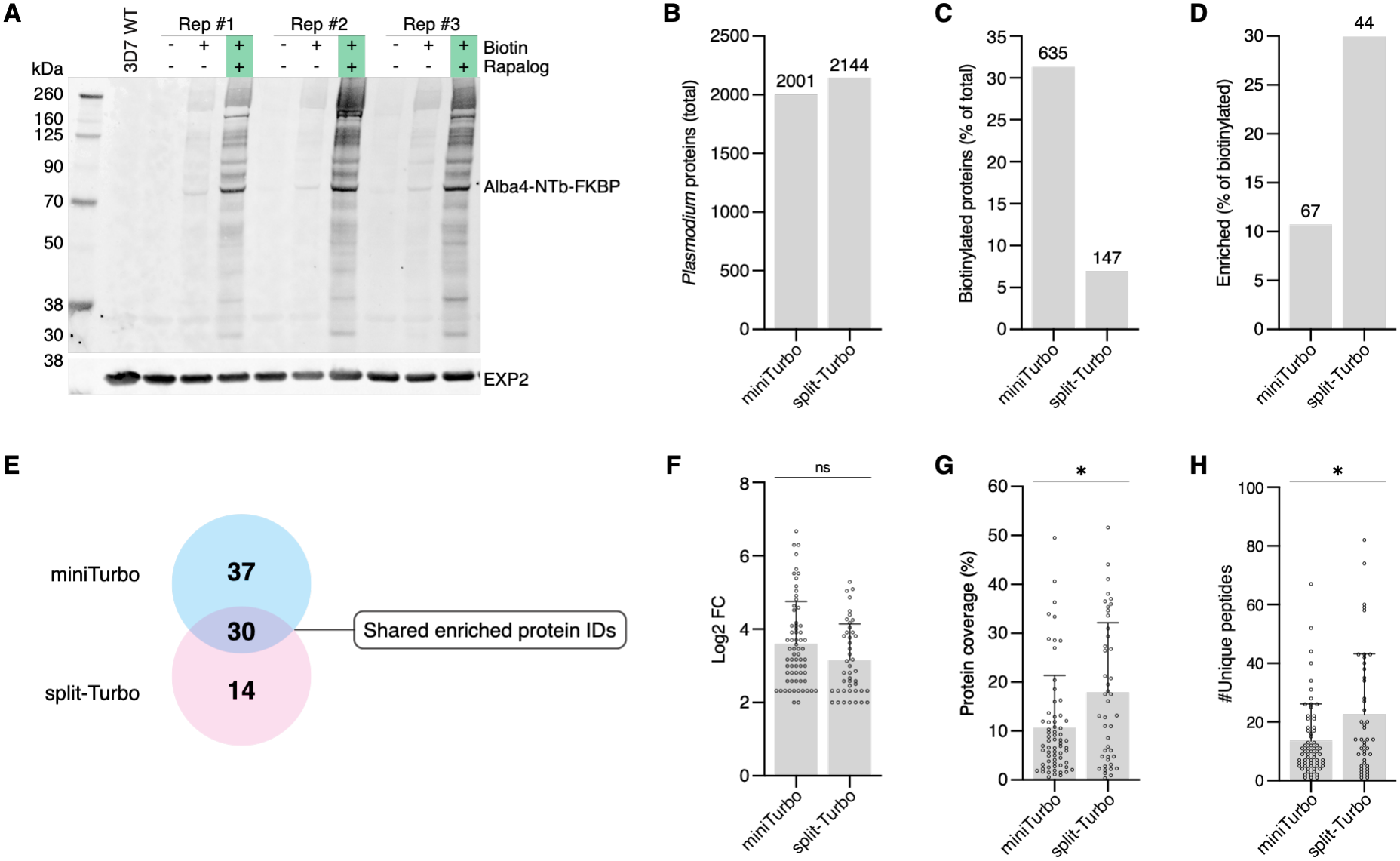
Comparative analysis of PfAlba4 split-TurboID and miniTurboID proximity labelling experiments. **A)** Western blot validation of late schizont stage PfAlba4-sTID harvests. Membranes were probed with streptavidin to detect biotinylated proteins. Shown are untreated (-/-), +Biotin only (+/-) and +Biotin/+Rapalog (+/+) treated samples, bands indicate biotinylation. A band corresponding to the expected size of PfAlba4-NTb-FKBP (∼73 kDa) is indicated and is more prominent in (+/+) lanes. Anti-EXP2 and 3D7 WT lysate were used as loading controls. Replicates #1-#3 are shown. **B)** Total number of *Plasmodium* proteins identified in previous PfAlba4-mTID data (n=4) (Zadow et al, manuscript submitted) and in this PfAlba4-sTID data (n=5), **C)** the percentage of these proteins that were biotinylated in PfAlba4-mTID and PfAlba4-sTID and **D)** the proportion of these protein IDs that were significantly enriched. Significance for split-TurboID experiments was determined as proteins with biotinylated peptides in the +Biotin/+Rapalog samples only and a minimum log2 fold-change (FC) of 2. Where proteins were also found biotinylated in the +Biotin only control, an additional threshold was added to only include those that appeared in a minimum of 3 +Biotin/+Rapalog replicates. Number of proteins is indicated above each graph. **E)** Previous miniTurboID experiments identified a total of 67 enriched PfAlba4 proximal proteins. PfAlba4 split-TurboID identified 44 enriched proteins and 30 of these proteins (∼68%) overlapped with the PfAlba4 miniTurboID list. **F)** Comparison of log2 foldchange, **G)** protein coverage and, **H)** the number of unique peptides assigned to each enriched protein in miniTurboID and split-TurboID data. Log2FC = fold change in the number of biotinylated peptides for a given protein in +Biotin/+Rapalog versus +Biotin only control samples; protein coverage = percentage of protein sequence with corresponding peptides; unique peptides = number of biotinylated peptide sequences assigned to this protein only. Mann-Whitney was used for all statistical analysis, individual data points are represented by dots, and graph columns indicate the mean for each dataset. Error bars show the standard deviation.

Mass spectrometric analysis of PfAlba4-sTID harvests identified a total of 2,144 *P. falciparum* proteins, similar in number to previous PfAlba4-mTID experiments, which identified 2,001 *P. falciparum* proteins (Figure 4b; Supplementary data 3). Analysis of biotinylated proteins revealed that a much higher percentage of proteins were biotinylated in the PfAlba4-mTID experiments compared to PfAlba4-sTID (31.4% and 6.9% of total protein IDs respectively) (Figure 4c). However, this comparison of the total number of biotinylated proteins also includes those that are biotinylated non-specifically. To get a better sense of the efficiency and specificity of the split-TurboID system, we compared the number of biotinylated proteins significantly enriched in the miniTurboID and the split-TurboID datasets (67 mTID: 44 sTID) and their relative proportions of total biotinylated proteins (10.7% mTID: 29.9% sTID). This indicated that while split-TurboID identified a lower number of significantly enriched proteins, a larger proportion of the total biotinylated proteins were found to meet the significance threshold in split-TurboID experiments, likely due to reduced background biotinylation (Figure 4c, d). Proteins were considered to be significantly enriched if they had a biotinylated peptide log2 fold change ≥2 in the +Biotin/+Rapalog samples compared to the +Biotin only controls and A) appeared in at least two +Biotin/+Rapalog replicates and were absent in all control samples or B) were present in control samples but appeared in at least three +Biotin/+Rapalog replicates.

There were 44 significantly enriched proteins identified for PfAlba4-sTID (Supplementary data 3) and 30 of these were found to overlap with the 67 significantly enriched proteins identified in the PfAlba4-mTID experiments (Figure 4e, Table 1). Comparison of high confidence proteins from both data showed that there was no significant difference between the log2 fold change of biotinylated peptides between the miniTurboID and split-TurboID experiments (Figure 4f). A comparison of protein sequence coverage and the number of unique peptide sequences identified for enriched proteins was observed to be higher using split-TurboID (Figure 4g, h), possibly due to the differences in labelling time (30-minutes for miniTurboID experiments versus 4-hours for split-TurboID). This longer labelling window would allow more opportunities for the BirA* to access less frequently exposed lysine residues for biotinylation of interacting proteins.

**Table 1.** PfAlba4 Split-TurboID and mini-TurboID overlapping enriched protein IDs.

| Accession # | Description | Split-TurboID |  | mini-TurboID |  |
|---|---|---|---|---|---|
|  |  | log2FC | P-value | log2FC | P-value |
| PF3D7_0108300 | conserved Plasmodium protein, unknown function | 5.29 | 0.00 | 5.17 | 0.00 |
| PF3D7_1415400 | Btz domain-containing protein, putative | 5.09 | 0.06 | 3.70 | 0.29 |
| PF3D7_1445700 | conserved Plasmodium protein, unknown function | 5.04 | 0.03 | 3.32 | 0.20 |
| PF3D7_1110400 | RNA-binding protein, putative | 4.49 | 0.00 | 5.52 | 0.00 |
| PF3D7_0511500 | RNA pseudouridylate synthase, putative | 4.39 | 0.00 | 6.30 | 0.01 |
| PF3D7_0604500 | GYF domain-containing protein, putative | 4.27 | 0.00 | 6.04 | 0.00 |
| PF3D7_0703500 | erythrocyte membrane-associated antigen | 4.25 | 0.00 | 4.09 | 0.18 |
| PF3D7_0706500 | conserved Plasmodium protein, unknown function | 4.04 | 0.00 | 3.91 | 0.00 |
| PF3D7_1021900 | PHAX domain-containing protein, putative | 3.91 | 0.07 | 4.09 | 0.31 |
| PF3D7_1224300 | polyadenylate-binding protein 1, putative | 3.91 | 0.00 | 3.46 | 0.11 |
| PF3D7_1202600 | nuclear export mediator factor NEMF, putative | 3.81 | 0.10 | 3.00 | 0.28 |
| PF3D7_1209000 | mago-binding protein, putative | 3.81 | 0.00 | 2.32 | 0.28 |
| PF3D7_1107300 | polyadenylate-binding protein-interacting protein 1, putative | 3.77 | 0.00 | 5.52 | 0.00 |
| PF3D7_1036900 | conserved Plasmodium protein, unknown function | 3.62 | 0.00 | 6.67 | 0.00 |
| PF3D7_1417200 | CCR4-NOT transcription complex subunit NOT1, putative | 3.46 | 0.00 | 3.46 | 0.24 |
| PF3D7_1145400 | dynamin-like protein | 3.32 | 0.00 | 3.17 | 0.15 |
| PF3D7_1365000 | conserved Plasmodium protein, unknown function | 3.17 | 0.01 | 2.32 | 0.18 |
| PF3D7_1423700 | NTF2-like domain-containing protein, putative | 3.13 | 0.00 | 4.64 | 0.20 |
| PF3D7_0811300 | CCR4-associated factor 1 | 3.00 | 0.06 | 3.70 | 0.25 |
| PF3D7_1366900 | conserved protein, unknown function | 2.86 | 0.00 | 3.70 | 0.27 |
| PF3D7_1019000 | eukaryotic translation initiation factor subunit eIF2A, putative | 2.66 | 0.01 | 4.91 | 0.00 |
| PF3D7_1308900 | mRNA-decapping enzyme 2, putative | 2.58 | 0.18 | 3.17 | 0.10 |
| PF3D7_1426100 | transcription factor BTF3, putative | 2.58 | 0.03 | 3.17 | 0.16 |
| PF3D7_0607000 | translation initiation factor IF-2, putative | 2.50 | 0.07 | 2.32 | 0.13 |
| PF3D7_0606600 | WD repeat-containing protein, putative | 2.46 | 0.02 | 5.04 | 0.00 |
| PF3D7_1004200 | WD repeat-containing protein, putative | 2.32 | 0.15 | 2.58 | 0.14 |
| PF3D7_0510100 | KH domain-containing protein, putative | 2.00 | 0.27 | 6.30 | 0.01 |
| PF3D7_1231800 | asparagine-rich protein, putative | 2.00 | 0.02 | 4.00 | 0.26 |
| PF3D7_0704600 | HECT-type E3 ubiquitin ligase UT | 2.00 | 0.18 | 3.46 | 0.30 |
| PF3D7_0212100 | conserved Plasmodium protein, unknown function | 2.00 | 0.27 | 2.32 | 0.30 |

**log2FC:** fold change in the number of biotinylated peptides detected in the +Biotin/+Rapalog samples versus the +Biotin only control; **p-value:** harmonic mean p-value derived from biotinylated peptides.

Due to the difference in the number of biotinylated proteins detected between PfAlba4-sTID (147) and PfAlba4-mTID (635) experiments, we compared the lists of total biotinylated proteins, inclusive of non-significant protein IDs, to look for any differences in biotinylated proteins and functions within a specific experiment (Supplementary data 3). Of the 635 biotinylated proteins identified in the PfAlba4-mTID data, 533 of these were unique to this dataset and GO analysis indicated enrichments for nuclear proteins and processes. For the 45 proteins biotinylated exclusively in PfAlba4-sTID experiments only two enrichments were observed and these were for pellicle/inner membrane pellicle complex proteins. Microscopy analysis of the miniTurboID episome used for PfAlba4 showed that in the absence of rapalog, this episome partially localised to the nucleus (Zadow *et al*, *Manuscript submitted*). Due to the constitutive activity of the biotin ligase carried on the miniTurboID episome, this could explain the background enrichment of nuclear proteins observed in these data.

There were 102 proteins found to be biotinylated across both the miniTurbo and split-Turbo datasets and GO enrichments mirrored those of the individual experiments and included CCR4-NOT complex proteins, and proteins involved in negative regulatory processes (Supplementary figure 6c-e). Overall, these results indicate that split-TurboID identifies potential protein interacting partners with high-specificity and high confidence, providing a robust and sensitive system for proximity labelling that can be better tolerated by the cell than we have found for miniTurboID.

### Split-TurboID facilitates identification of a high confidence core proxiome for PfAlba2

Following successful validation of split-TurboID using PfAlba4, we next generated a PfAlba2-sTID parasite line in the same manner. Tag integration was once again confirmed by PCR (Figure 5a) and expression of both PfAlba2-NTb-FKBP-GFP and CTb-FRB-3xHA was confirmed by western blot (Figure 5b, c). Immunofluorescence microscopy demonstrated the same nucleocytoplasmic localisation for PfAlba2 as was previously observed (Figure 5d; Supplementary figure 7a) and a cytoplasmic localisation was observed for the CTb-FRB-HA component, similar to PfAlba4-sTID (Figure 5e; Supplementary figure 7b). Biotinylation efficiency was also tested at 1-, 2-, and 4-hours incubation and mirrored the previous PfAlba4-sTID results, demonstrating specific enrichment in +Biotin/+Rapalog samples and increasing protein biotinylation to 4 hours (Supplementary figure 7c-f). Dimerisation-induced biotinylation in the presence of Rapalog for the PfAlba2-sTID line was also verified by microscopy (Supplementary figure 8a). Late-stage PfAlba2-sTID parasites were then harvested for proteomics analysis, with efficient biotinylation confirmed by western blot prior to downstream processing (Figure 5f; Supplementary figure 8b, c).

**Figure 5.**
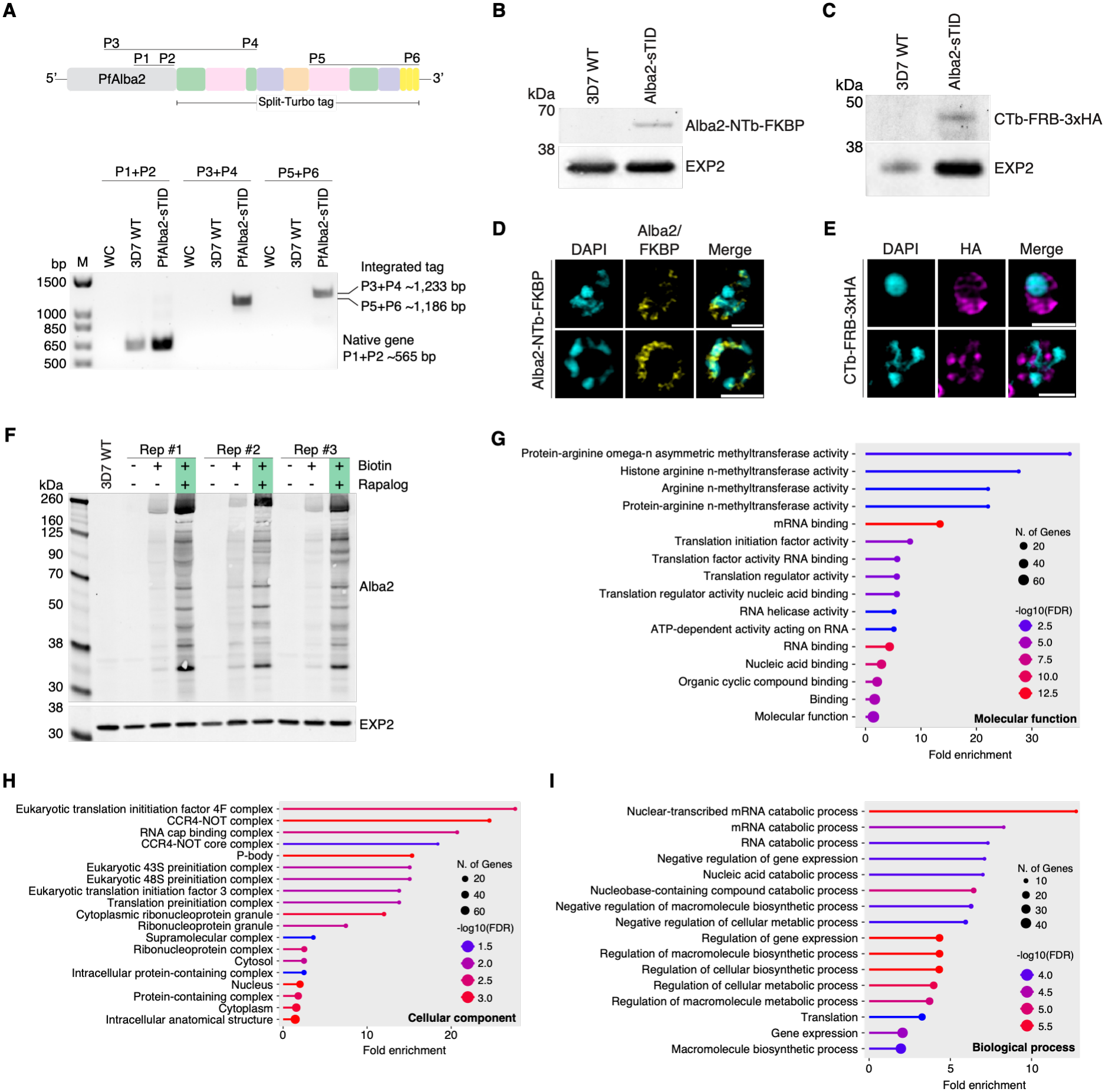
Identification of a high confidence PfAlba2 proxiome using split-TurboID. **A)** Successful endogenous tagging of PfAlba2 was confirmed by PCR. P1 and P2 target a region of the native gene and a band at the expected size of ∼565 bp was observed in both the PfAlba2-sTID gDNA and the 3D7 WT control. P3 targets a region upstream of the 3’ homology region used in the transfection plasmid and P4 targets the N-terminal BirA* fragment. Presence of the expected ∼1,233 bp band in the PfAlba2-sTID lane indicates successful integration, no band was present in the 3D7 WT control. P5 and P6 indicate the presence of the C-terminal portion of the split-TurboID tag, no band is observed in 3D7 WT as expected while a band of ∼1,186 bp is present in the PfAlba2-sTID lane. Bands were absent from water controls (WC) for all primers pairs. **B)** Western blots confirmed expression of PfAlba2 fused with the N-terminal fragment and FKBP (Alba2-NTb-FKBP) with the corresponding band detected with anti-FKBP at ∼61 kDa). Anti-EXP2 and 3D7 WT were used as loading controls. **C)** Western blots probed with anti-HA confirmed expression of the C-terminal BirA* fragment (CTb-FRB-3xHA) at the expected size of ∼45 kDa and anti-EXP2 and 3D7 WT were used as loading controls. **D)** Super-resolution microscopy confirmed the localisation of PfAlba2-NTb-FKBP (yellow) which was consistent with prior observations of a largely cytosolic localisation. Nuclei stained with DAPI (cyan) and, **E)** localisation of the CTb-FRB-3xHA tag using anti-HA antibody (magenta). Scale bars = 2 μm. **F)** Western blot validation of PfAlba2-sTID late schizont stage parasite harvests. Samples from replicates #1 - #3 are shown (n=4) and bands correspond to biotinylated proteins detected by streptavidin antibody. Shown are the untreated control (-/-), +Biotin only control (+/-) and the +Biotin/+Rapalog treated samples (+/+). A band corresponding to the expected size of PfAlba2-NTb-FKBP is also indicated (∼61 kDa) and 3D7 WT lysate and anti-EXP2 were used as loading controls. **G)** Gene Ontology analysis of the PfAlba2 high confidence proxiome highlighted enriched molecular functions, **H)** cellular components and, **I)** Biological processes in these data. Analysis was performed using default parameters and terms are sorted by fold enrichment with number of genes and statistical significance indicated in the associated legends of each graph (ShinyGO 0.85).

Proteomics analysis identified 101 significantly enriched potential interacting and proximal proteins for PfAlba2 using the same enrichment criteria as for PfAlba4-sTID (Table 2; Supplementary data 4). Of these proteins, 55 appeared across all four replicates and 54 were found only in the +Biotin/+Rapalog samples. Additionally, ∼12% of the enriched proteins are currently annotated as conserved with unknown function. Proteins involved in translation initiation were highly enriched including putative eukaryotic translation initiation factor (eIF) subunit 2A (PF3D7_1019000), eIF4E and eIF4G (PF3D7_0111800, PF3D7_1312900), and putative eIF3 subunits A, D, and E (PF3D7_1212700, PF3D7_1007900, PF3D7_0528200) (Figure 5g-i, figure 6). Multiple ATP-dependent RNA helicases were also identified including DDX6 (PF3D7_0320800) and the putative DBP1, DBP10, and DHX57 (PF3D7_0810600, PF3D7_0827000, PF3D7_0310500). Enrichments for methyltransferases are also notable and included protein-arginine N-methyltransferase 1 (PRMT1; PF3D7_1426200) and the putative histone-arginine methyltransferase CARM1 (PF3D7_0811500).

**Figure 6.**
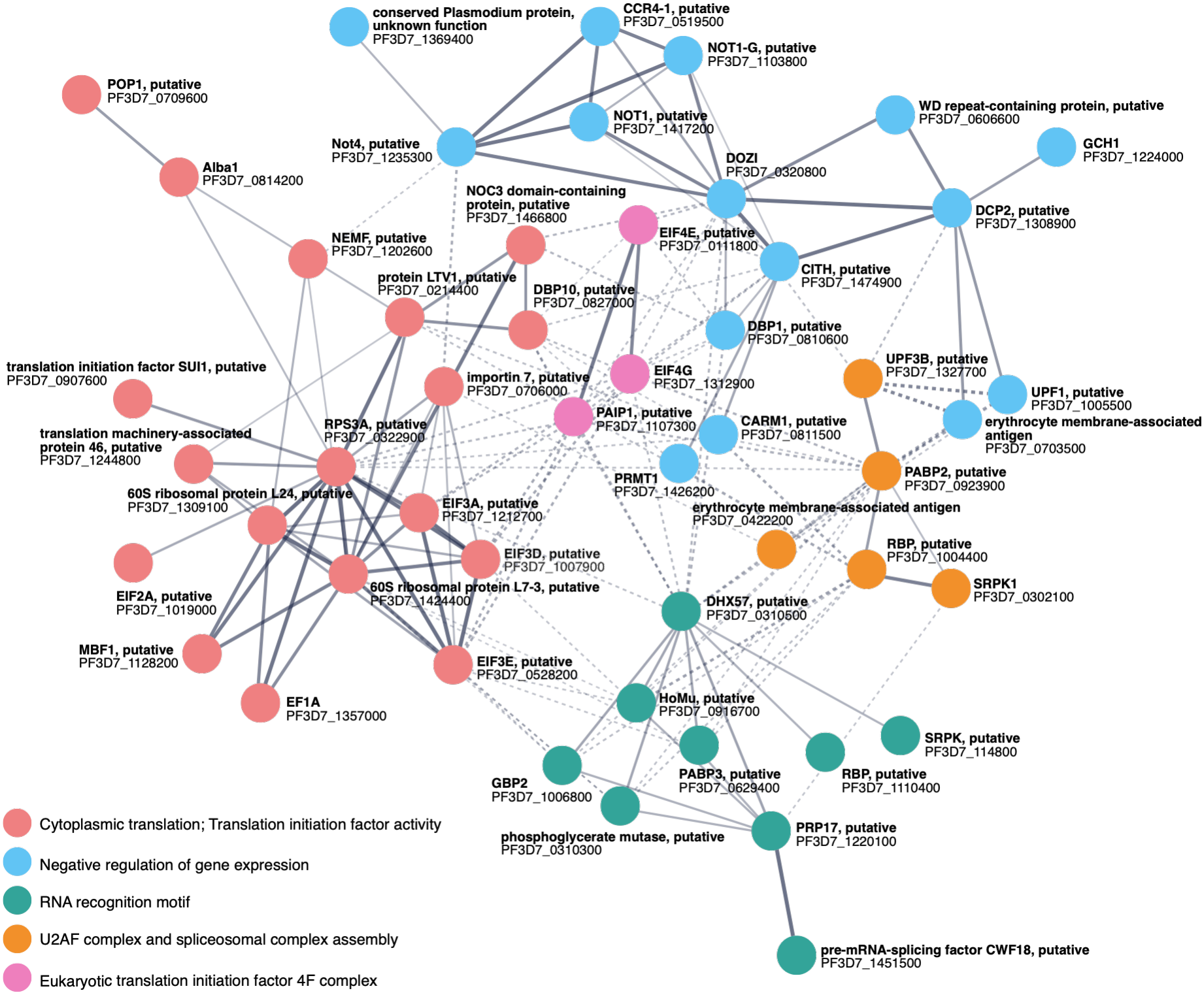
Network analysis of the PfAlba2 proxiome highlights enrichment of proteins involved in key cell regulatory processes. Network analysis was undertaken in StringDB using K-means clustering. The top five clusters are represented and include cytoplasmic translation and translation initiation factor activity (red); negative regulation of gene expression (blue); RNA recognition motif (green); U2AF complex and spliceosomal complex assembly (orange) and; Eukaryotic translation initiation factor 4F complex (pink).

**Table 2.**
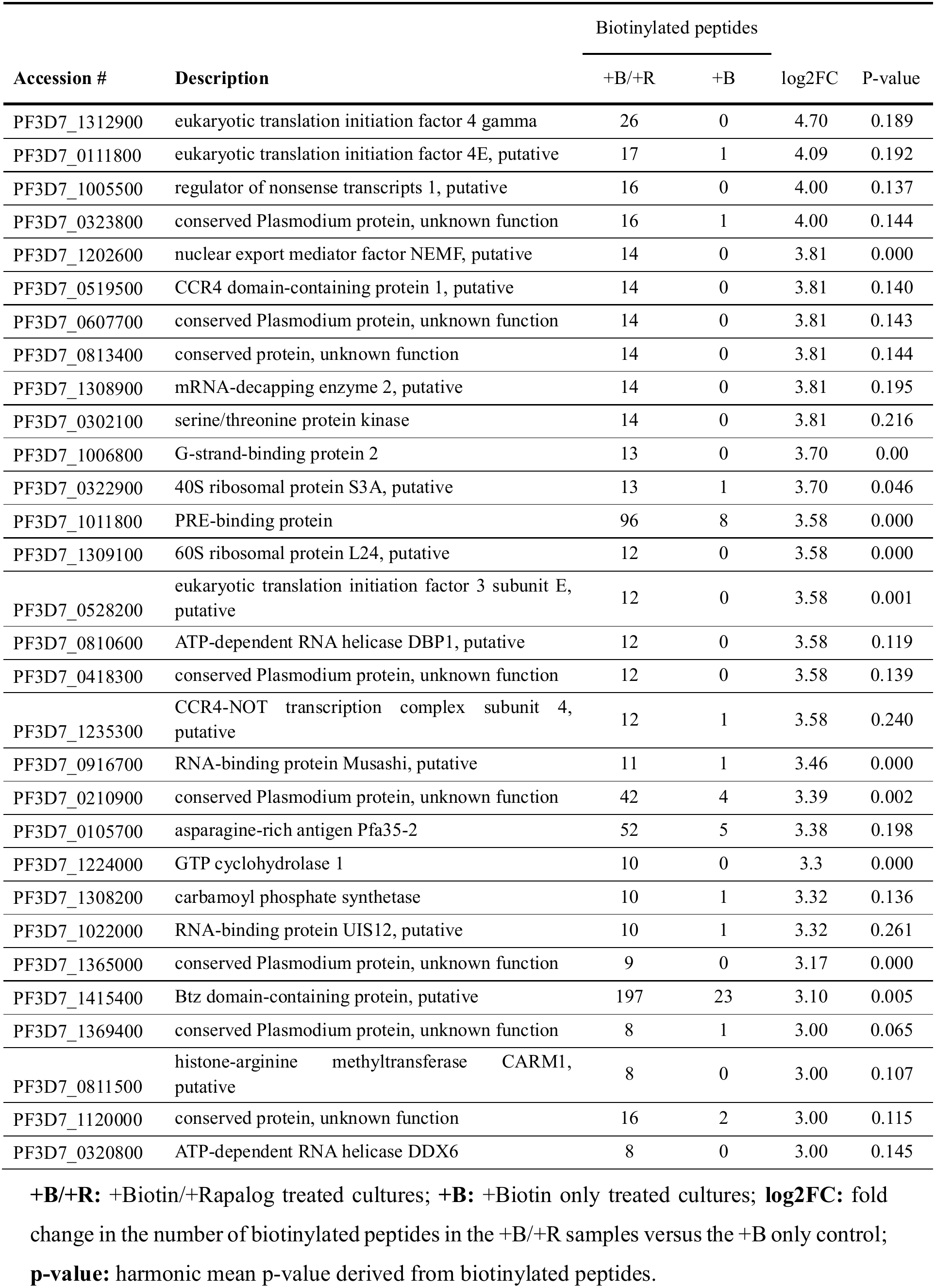
PfAlba2 top 30 enriched proteins identified with split-TurboID.

|  |  | Biotinylated peptides |  |  |  |
|---|---|---|---|---|---|
| **Accession #** | **Description** | +B/+R | +B | log2FC | P-value |
| PF3D7_1312900 | eukaryotic translation initiation factor 4 gamma | 26 | 0 | 4.70 | 0.189 |
| PF3D7_0111800 | eukaryotic translation initiation factor 4E, putative | 17 | 1 | 4.09 | 0.192 |
| PF3D7_1005500 | regulator of nonsense transcripts 1, putative | 16 | 0 | 4.00 | 0.137 |
| PF3D7_0323800 | conserved Plasmodium protein, unknown function | 16 | 1 | 4.00 | 0.144 |
| PF3D7_1202600 | nuclear export mediator factor NEMF, putative | 14 | 0 | 3.81 | 0.000 |
| PF3D7_0519500 | CCR4 domain-containing protein 1, putative | 14 | 0 | 3.81 | 0.140 |
| PF3D7_0607700 | conserved Plasmodium protein, unknown function | 14 | 0 | 3.81 | 0.143 |
| PF3D7_0813400 | conserved protein, unknown function | 14 | 0 | 3.81 | 0.144 |
| PF3D7_1308900 | mRNA-decapping enzyme 2, putative | 14 | 0 | 3.81 | 0.195 |
| PF3D7_0302100 | serine/threonine protein kinase | 14 | 0 | 3.81 | 0.216 |
| PF3D7_1006800 | G-strand-binding protein 2 | 13 | 0 | 3.70 | 0.00 |
| PF3D7_0322900 | 40S ribosomal protein S3A, putative | 13 | 1 | 3.70 | 0.046 |
| PF3D7_1011800 | PRE-binding protein | 96 | 8 | 3.58 | 0.000 |
| PF3D7_1309100 | 60S ribosomal protein L24, putative | 12 | 0 | 3.58 | 0.000 |
| PF3D7_0528200 | eukaryotic translation initiation factor 3 subunit E, putative | 12 | 0 | 3.58 | 0.001 |
| PF3D7_0810600 | ATP-dependent RNA helicase DBP1, putative | 12 | 0 | 3.58 | 0.119 |
| PF3D7_0418300 | conserved Plasmodium protein, unknown function | 12 | 0 | 3.58 | 0.139 |
| PF3D7_1235300 | CCR4-NOT transcription complex subunit 4, putative | 12 | 1 | 3.58 | 0.240 |
| PF3D7_0916700 | RNA-binding protein Musashi, putative | 11 | 1 | 3.46 | 0.000 |
| PF3D7_0210900 | conserved Plasmodium protein, unknown function | 42 | 4 | 3.39 | 0.002 |
| PF3D7_0105700 | asparagine-rich antigen Pfa35-2 | 52 | 5 | 3.38 | 0.198 |
| PF3D7_1224000 | GTP cyclohydrolase 1 | 10 | 0 | 3.3 | 0.000 |
| PF3D7_1308200 | carbamoyl phosphate synthetase | 10 | 1 | 3.32 | 0.136 |
| PF3D7_1022000 | RNA-binding protein UIS12, putative | 10 | 1 | 3.32 | 0.261 |
| PF3D7_1365000 | conserved Plasmodium protein, unknown function | 9 | 0 | 3.17 | 0.000 |
| PF3D7_1415400 | Btz domain-containing protein, putative | 197 | 23 | 3.10 | 0.005 |
| PF3D7_1369400 | conserved Plasmodium protein, unknown function | 8 | 1 | 3.00 | 0.065 |
| PF3D7_0811500 | histone-arginine methyltransferase CARM1, putative | 8 | 0 | 3.00 | 0.107 |
| PF3D7_1120000 | conserved protein, unknown function | 16 | 2 | 3.00 | 0.115 |
| PF3D7_0320800 | ATP-dependent RNA helicase DDX6 | 8 | 0 | 3.00 | 0.145 |

**+B/+R:** +Biotin/+Rapalog treated cultures; **+B:** +Biotin only treated cultures; **log2FC:** fold change in the number of biotinylated peptides in the +B/+R samples versus the +B only control; **p-value:** harmonic mean p-value derived from biotinylated peptides.

The PfAlba2 proxiome also contained several components of putative CCR4-NOT complex proteins including CCR4-1, NOT4, NOT1-G, and NOT1 (PF3D7_0519500, PF3D7_1235300, PF3D7_1103800, PF3D7_1417200) (Figure 5g-i, figure 6; Supplementary data 4). Network analysis in StringDB further highlighted enrichments for proteins containing RNA recognition motifs and proteins involved in mRNA processing (Figure 6).

Notably, two other PfAlba proteins were also identified with high confidence, PfAlba1 (PF3D7_0814200) and PfAlba4 (PF3D7_1347500). Enrichment of these proteins could indicate heterodimerisation between these Alba proteins, as has been observed for some Alba proteins in other model organisms (Hands-Taylor *et al*, 2010; Jelinska *et al*, 2005; Welting *et al*, 2007). Potential interactions between PfAlba1 and PfAlba4 have recently been indicated in a study by Pazicky *et al* (2025) however this remains to be experimentally confirmed.

### RIP-Seq analysis implicated PfAlba2 in translational repression of pathways including the ribosome and RNA processing

Given the reported roles of Alba proteins in regulating RNA and the associations between PfAlba2 and RNA regulatory proteins identified in the split-TurboID PfAlba2 data, we next sought to identify what RNA transcripts might be interacting with PfAlba2 by RIP-seq. RIP-seq identified 380 significantly enriched (≥2.5-fold enrichment and FDR ≤0.05) PfAlba2-associated transcripts from 5,043 transcripts evaluated in schizont-stage parasites (Figure 7a; Supplementary data 5), with representative transcripts reflecting varying degrees of enrichment following PfAlba2 immunoprecipitation (Figure 7b). To identify putative cis-regulatory elements mediating PfAlba2 target recognition, we performed discriminative motif analysis (XSTREME) on the 3’UTRs of PfAlba2 RIP-Seq targets against non-target *P. falciparum* 3’UTRs as background. STREME identified a 15-nucleotide poly-U/GAAU element (AUUUUUUUUUUGAAU) (Figure 7c) enriched 12.3-fold in target sequences relative to background (3.64% vs 0.29%; SEA p = 4.43 × 10⁻⁷ q = 4.22 × 10⁻⁵). Tomtom comparison against the Ray2013 RNAcompete RBP motif database revealed significant similarity to the binding profiles of the Drosophila translational repressor Sex-lethal (SXL) and multiple ELAV/Hu-family proteins, all of which recognise poly-U tracts in 3’UTRs to repress translation (Supplementary data 6) (Ray *et al*, 2013b). Notably, FIMO scanning identified this element in the 3’UTR of PfAlba1 mRNA itself, a potential direct heterodimeric partner of PfAlba2, suggesting a potential auto-regulatory circuit governing Alba complex stoichiometry (Supplementary data 6). These findings indicate that a subset of PfAlba2 targets are distinguished by a poly-U/GAAU element analogous to ARE-like sequences bound by translational repressors in other eukaryotes, consistent with the proposed functional parallel between the Plasmodium DOZI/CITH/Alba machinery and the mammalian ARE-mediated mRNA storage system (Min *et al*, 2024; Rios *et al*, 2025; Wu & Brewer, 2012).

**Figure 7.**
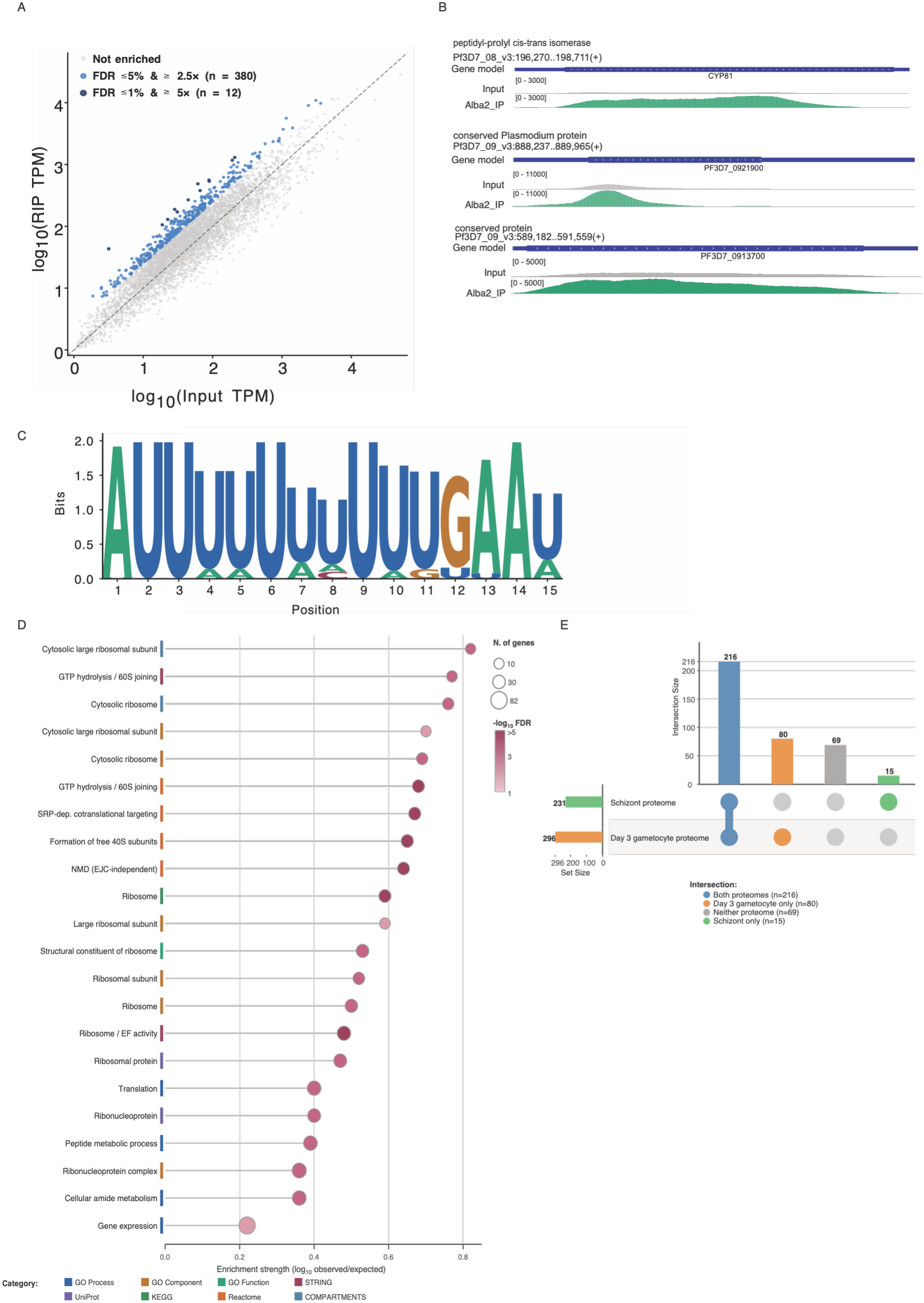
RIP-Seq identifies 380 PfAlba2-associated mRNA targets in *P. falciparum* schizonts. **A)** Scatter plot comparing log₁₀-transformed TPM values between the PfAlba2 RIP pulldown (RIP TPM, y-axis) and total RNA input (Input TPM, x-axis) for all genes detected in the RIP-Seq experiment. Grey points represent non-enriched transcripts. Blue points represent primary targets enriched at FDR ≤ 5% and ≥ 2.5-fold (n = 380), with dark blue points indicating high-confidence targets at FDR ≤ 1% and ≥ 5-fold enrichment (n = 12). The dashed line indicates equal abundance between pulldown and input. **B)** Representative IGV genome browser tracks showing PfAlba2 RIP-Seq enrichment over three example target genes. Tracks display the gene model (top), total RNA input (middle, grey), and Alba2 immunoprecipitate (bottom, green) for peptidyl-prolyl cis-trans isomerase (PF3D7_0803000; rank 3, 6.7×), a conserved Plasmodium protein (PF3D7_0921900), and a conserved protein (PF3D7_0913700). Read depth scales are indicated in brackets. **C)** Sequence logo of the top-ranked STREME-1 motif (AUUUUUUUUUUGAAU) identified by discriminative XSTREME analysis of PfAlba2 target 3’UTRs against 3,777 non-target *P. falciparum* 3’UTRs as background (MEME Suite v5.5). Letter height represents information content in bits (maximum 2 bits). The motif was enriched 12.3-fold in target sequences relative to background (3.64% vs 0.29%; SEA p = 4.43 × 10⁻⁷, E-value = 1.12 × 10⁻⁴, q = 4.22 × 10⁻⁵) and shows significant Tomtom similarity to the binding profiles of the Drosophila translational repressor Sex-lethal (SXL) and multiple ELAV/Hu-family proteins. **D)** Horizontal lollipop plot showing functional enrichment of PfAlba2 RIP-Seq targets. The x-axis shows enrichment strength (log₁₀ observed/expected gene count). Dot size is proportional to the number of observed genes and dot colour indicates −log₁₀ FDR (light pink: FDR ∼0.04; dark red: FDR < 0.001). Categories are colour-coded as indicated. Enrichment analysis was performed using STRING (v11.5) against a background of all *P. falciparum* 3D7 genes. **E)** UpSet plot comparing PfAlba2 RIP-Seq targets (n = 380) against stage-specific proteomes from *P. falciparum* schizonts and wild-type Day 3 gametocytes. The upper bar chart shows intersection sizes; the dot matrix below indicates which proteome sets are active in each intersection. Horizontal bars on the left show total set sizes. 216 targets (56.8%) were detected at the protein level in both stages, 80 (21.1%) exclusively in Day 3 gametocytes, 15 (3.9%) exclusively in schizonts, and 69 (18.2%) were undetected in either proteome.

Using an FDR <0.05 threshold, functional enrichment analysis showed that PfAlba2-associated transcripts were enriched for translation, peptide and amide metabolism, gene expression, structural constituents of the ribosome, cytosolic ribosomes and ribonucleoprotein complexes (Figure 7d; Supplementary data 7). KEGG and Reactome analyses further highlighted ribosome, co-translational protein targeting, ribosomal subunit assembly, nonsense-mediated mRNA decay, RNA metabolism and rRNA processing (Figure 7d; Supplementary data 7). Together, these analyses suggest translation, ribosome function and RNA processing as the dominant features of the PfAlba2 target set. Significant PfAlba2 targets included multiple cytosolic ribosomal proteins (L26, PF3D7_0312800; L39, PF3D7_0611700; L35, PF3D7_1124900; L7, PF3D7_0307200; S17, PF3D7_1242700; S26, PF3D7_0217800), rRNA-processing factors, the RNA helicase DDX41 (PF3D7_0527900), mRNA-capping machinery and PfAlba1 (PF3D7_0814200) itself. Other notable targets included calcineurin, MAP kinase 1 (PF3D7_1431500), apicoplast and mitochondrial ribosomal components, origin recognition complex (ORC) subunits, and schizont stage-associated proteins including RON4 (PF3D7_1116000), RH1 (PF3D7_0402300), RH5 (PF3D7_0424100), and HAP2P (PF3D7_0816300).

During blood stage development, a subset of parasites will differentiate into male and female gametocytes which are essential for the continued transmission of these parasites. Previous studies have implicated Plasmodium Alba proteins in the suppression of male gametocytes in particular so to investigate this further we compared the enriched PfAlba2-associated transcripts with schizont and day 3 gametocyte proteomes (Mair *et al*, 2006; Muñoz *et al*., 2017; Rios *et al*., 2025). Analysis against stage-specific proteomes showed that 231 of the 380 PfAlba2-associated transcripts (60.8%) had corresponding protein evidence in schizonts, leaving 149 without detectable protein products in this stage (Figure 7e; Supplementary data 8) (Chisholm *et al*, 2026; Marapana *et al*, 2026). Of these 149 transcripts, 80 (53.7%) were detected at the protein level in wild-type Day 3 gametocytes. Consequently, 311 of 380 PfAlba2 targets (81.8%) had protein-level evidence across the schizont and Day 3 gametocyte datasets, while 69 remained undetected in either proteome (Figure 7e; Supplementary data 8). This demonstrates substantial stage-dependent variation in the protein detection of PfAlba2-associated transcripts.

## Discussion

The functional characterisation of *Plasmodium* spp. Alba proteins has been limited to date, particularly in the important human-infecting species *P. falciparum*, where difficulties in generating endogenously tagged lines have restricted the direct investigation of their biological roles. Here, we successfully generated two different endogenously tagged PfAlba2 transgenic lines, enabling investigation of its localisation, functional characterisation, and its protein and mRNA associations during late blood stage development. Our findings reveal associations between PfAlba2, translation-related proteins, and transcripts involved in RNA metabolism and erythrocyte invasion, providing new insight into its potential role in post-transcriptional regulation.

Endogenous tagging confirmed that PfAlba2 is ubiquitously expressed and largely cytoplasmic during blood stage development, which is consistent with previously reported studies that utilised in-house antibodies or ectopic overexpression systems (Acharya *et al*., 2025; Chêne *et al*., 2012). Conditional mislocalisation of PfAlba2 resulted in a modest impact on parasite growth, with a subset of parasites displaying aberrant morphologies that included stress vacuoles, cellular disorganisation, and membrane irregularities. These findings suggest that PfAlba2 contributes to normal *P. falciparum* asexual blood stage development, consistent with its predicted essentiality in genome-wide saturation mutagenesis studies (Zhang *et al*., 2018). However, the relatively modest growth defect observed following conditional mislocalisation may reflect incomplete functional disruption and therefore does not exclude an essential role for PfAlba2.

A novel contribution of this study was the establishment of a customised split-TurboID system for proximity labelling in *P. falciparum.* Repeat attempts to generate a PfAlba2 miniTurboID system were unsuccessful and suggested that the episomal expression of the miniTurboID was not well tolerated in the PfAlba2^2xFKBP-GFP^ background, a problem we and others have seen for different proteins (*personal communications*). This prompted the development of an alternative approach based on rapalog-inducible reconstitution of the biotin ligase. In establishing a *Plasmodium* compatible split-TurboID system, we sought to reduce the cellular toxicity sometimes seen with miniTurboID, improve BirA* accessibility, provide accessible and flexible tagging options, enable the long-term use of standard h-RPMI culture medium for regular parasite maintenance, and reduce the non-specific background biotinylation typically observed with miniTurboID (Branon *et al*., 2018a; May *et al*, 2020).

To ensure that this system would be an accessible option to others in the community, we chose to utilise the existing and well-established selection-linked integration (SLI) strategy for endogenous tagging of target proteins (Birnbaum *et al*., 2017). This method offers a straight-forward endogenous tagging approach using relatively inexpensive and common reagents. Additionally, incorporation of a T2A skip peptide between the NTb and CTb fragments facilitates generation of the required transgenic parasite lines in a single transfection step, and the separation of the NTb and CTb fragments addresses the issues of cellular toxicity sometimes seen with the highly active BirA* of miniTurboID.

The system also offers flexibility for future applications, with additional restriction sites incorporated to facilitate modifications for different target proteins. Although effective for the highly expressed PfAlba proteins, its dependence on target protein expression may limit labelling efficiency for less abundant proteins. Validation using PfAlba4 demonstrated inducible biotinylation and recovery of a substantial proportion of the high confidence proteins previously identified using miniTurboID. Although split-TurboID identified fewer significantly enriched proteins overall, a greater proportion of its biotinylated proteins met the significance threshold, consistent with potentially reduced background labelling. However, differences in labelling duration and experimental configuration prevent direct conclusions regarding the relative sensitivity or specificity of the two approaches. Nevertheless, these findings establish split-TurboID as a valuable additional proximity-labelling strategy for investigating protein associations in malaria parasites, particularly where conventional miniTurboID approaches prove challenging.

Application of split-TurboID to PfAlba2 revealed a network of proximal proteins enriched for factors involved in translational initiation, including components of the eukaryotic translation initiation factor 4F (eIF4F) complex. In model organisms, the eIF4F complex facilitates the recruitment of ribosomes to mRNA (Gingras *et al*, 1999). It forms a heterotrimeric complex consisting of eIF4E (cap-binding; PF3D7_0111800), eIF4A (helicase; PF3D7_1468700), and eIF4G (bridging/scaffold, mediates interactions with polyadenylate-binding proteins; PF3D7_1312900) (Gingras *et al*., 1999). Both eIF4E and eIF4G were identified with high confidence in the PfAlba2 proxiome and interestingly, sequence analysis using the ELM database indicated that PfAlba2 possesses an eIF4E binding motif at its N-terminus (Kumar *et al*, 2020). In other organisms, eIF4G also contains an eIF4E binding domain as do eIF4E inhibitory proteins which compete with eIF4G for binding with eIF4E as a means of translational regulation (Richter & Sonenberg, 2005). These findings raise the possibility that PfAlba2 acts as an eIF4E inhibitory protein during late blood stage development, although direct binding and its potential effects on translational initiation remain to be investigated.

Other notable high confidence proteins identified in the PfAlba2 proxiome include the arginine methyltransferases PfPRMT1 and the putative PfCARM1, both are homologous to type I protein methyltransferases. ELM analysis of the PfAlba2 protein sequence identified a PRMT recognition motif (GGRGG) which is recognised by arginine methyltransferases and a sequence commonly found in RNA-binding proteins (Kumar *et al*., 2020). In other eukaryotes, protein-arginine methyltransferases play important roles in regulating RNA metabolism and translation (Bachand & Silver, 2004; Ferreira *et al*, 2020; Rodriguez, 2022; Sudhakar *et al*, 2024; Wolf, 2009). In *P. falciparum,* PfPRMT1 has been shown to methylate histone substrates including PfH4 and PfH2A as well as serine/arginine-rich splicing factor 1 (PF3D7_0517300), and PfPABP2 (PF3D7_1447000; also identified in PfAlba2-BioID) (Fan *et al*, 2009). Although PfCARM1 (also referred to as PRMT4) remains functionally uncharacterised, PlasmoDB annotations suggest that this may be an orthologue of the closely related PRMT3 (also a type I protein methyltransferase) based on homology with the *Toxoplasma gondii* protein TgPRMT3 (TGGT1_252420). Together, these findings suggest that arginine methylation may contribute to the regulation of PfAlba2-associated RNA regulatory processes, although direct methylation of PfAlba2 remains to be established.

The PfAlba4 and PfAlba2 proxiomes shared several functional enrichments, notably one of these was components of the CCR4-NOT complex. This highly conserved complex has broad regulatory functions in eukaryotes, including mRNA turnover, translational regulation and mRNA surveillance, although several of its components remain poorly characterised in *Plasmodium* spp. (Caulier *et al*, 2025; Collart, 2016). Given the involvement of CCR4-NOT in ribosome-associated quality control, its enrichment in our BioID data could potentially reflect an artefact arising from ribosome stalling associated with the T2A ribosomal skipping sequence incorporated into the split-TurboID construct. However, our data suggest that this is unlikely to be the sole explanation. In the PfAlba2 split-TurboID experiments, 43 biotinylated peptides corresponding to CCR4-NOT components were detected following combined biotin and rapalog treatment, compared with only four in the biotin-only control. Similarly, PfAlba4 split-TurboID identified 19 CCR4-NOT-associated biotinylated peptides in the induced condition and none in the biotin-only control. Importantly, CCR4-NOT components were also enriched in our previous PfAlba4 miniTurboID experiments, which lacked the T2A sequence, with 49 biotinylated peptides corresponding to CCR4-NOT components detected in the biotin and rapalog-treated samples and none in the biotin only controls (Zadow *et al*, *Manuscript submitted*). The consistent enrichment of CCR4-NOT components across these datasets, particularly in the absence of the T2A sequence, argues against an artefact arising solely from ribosomal skipping. Together these findings support a potential association between PfAlba proteins and CCR4-NOT-mediated post-transcriptional regulatory pathways.

RIP-seq analysis identified 380 PfAlba2-associated transcripts enriched in late blood stage development, including several encoding proteins involved in schizont egress and invasion. Notable transcripts include PfRALP1 (rhoptry-associated leucine zipper-like protein 1; PF3D7_0722200), PfRON4 (rhoptry neck protein 4; PF3D7_1116000), PfRH1 (reticulocyte-binding homologue 1; PF3D7_0402300), and PfRH5 (PF3D7_0424100). Interestingly, a previous study of PfAlba1 (PF3D7_0814200)-associated transcripts during trophozoite development also identified several invasion-related mRNAs, including PfRON4 (Vembar *et al*., 2015). That study further demonstrated that PfAlba1 negatively regulates the translation of four other schizont specific transcripts: PfRhopH3 (high molecular weight associated protein 3; PF3D7_0905400), PfCDPK1 (calcium-dependent protein kinase 1; PF3D7_0217500), PfRap1 (rhoptry-associated protein 1; PF3D7_1410400), and PfAMA1 (apical membrane antigen 1; PF3D7_1133400).

The overlap between previously reported PfAlba1-associated transcripts by Vembar *et al*. (2015) and our PfAlba2 RIP-seq dataset is particularly noteworthy as PfAlba1 was also identified as a high-confidence proximal protein in our PfAlba2 split-TurboID experiments. These findings suggest that PfAlba1 and PfAlba2 may participate in overlapping post-transcriptional regulatory pathways, potentially involving transcripts associated with erythrocyte invasion. PfAlba4 was also identified as a high-confidence proximal protein of PfAlba2, and the PfAlba2 and PfAlba4 proxiomes shared several proteins, further suggesting potential functional relationships among these family members. Together, these observations support the possibility that PfAlba proteins participate in shared RNA regulatory networks during asexual blood stage development, although their precise molecular interactions and regulatory functions remain to be established. It is notable that PfAlba2 was not reciprocally enriched in the PfAlba4 proxiome data however, this could potentially be due to tag location and a lack of access to biotinylation sites.

Alba proteins have been reported to form homo- and heteromeric assemblies in several organisms, with oligomerisation of PfAlba3 and PfAlba6 previously documented in *P. falciparum* (Banerjee *et al*, 2023; Chêne *et al*., 2012; Hands-Taylor *et al*., 2010; Jelinska *et al*., 2005; Nag *et al*, 2024; Welting *et al*., 2007). Previous co-immunoprecipitation studies have also suggested potential interactions between PfAlba family members, although these approaches cannot distinguish direct interactions from associations within larger protein compelxes (Muñoz *et al*., 2017; Vembar *et al*., 2015). Methods of recombinant protein expression have also been utilised and recent work by Pazicky *et al*. (2025), demonstrated heterodimerisation among recombinantly expressed *P. berghei* Alba proteins 1-3, as well as homodimerisation of PbAlba1, providing further evidence for Alba protein oligomerisation in malaria parasites.

Interestingly, the identification of PfAlba1 and PfAlba4 enrichments in the PfAlba2 proxiome suggests the possibility of heterodimer formation between these family members, so we explored this further by examining which regions of PfAlba1 and PfAlba4 were biotinylated in the PfAlba2 BioID. For PfAlba1, all detected biotinylated peptides were at the very N-terminus of the Alba domain and in a middle region of the protein overlapping with its distinctive Rpp25-like motif, with this same pattern observed for PfAlba4 (Supplementary figure 9a). While PfAlba1 and PfAlba4 were not enriched in the PfAlba4 split-TurboID data, it is notable that the few biotinylated peptides identified for these proteins in the PfAlba4 split-TurboID dataset were in the corresponding regions to those seen for these proteins in the PfAlba2 split-TurboID. Given that the biotinylating enzyme is located at the C-terminus of PfAlba2, we hypothesise that these Alba proteins were potentially interacting in a head-to-tail, back-to-back fashion similar to Alba-Alba interactions observed in other organisms, and this was further supported by the predicted interactions generated using AlphaFold (Supplementary figure 9b, c) (Goyal *et al*., 2016; Jumper *et al*, 2021). It is also notable that, due to the nature of the split-TurboID system, the two portions of the BirA* require close proximity of ∼2-5 nm to restore enzyme activity (Cho *et al*., 2020a). This results in a highly restricted biotinylation radius for this system and adds further confidence to predictions of true interacting proteins.

An additional observation from our RIP-seq analysis was the identification of PfAlba1 mRNA as a PfAlba2-associated transcript. Notably, the enriched 3’UTR motif identified among PfAlba2-associated transcripts was also present in the PfAlba1 3’ UTR, raising the possibility that PfAlba2 may contribute to the regulation of PfAlba1 expression. Regulatory interactions between Alba family members are not uncommon and have been reported in other apicomplexan parasites. In *T. gondii,* it has been demonstrated that TgAlba1 (TGME49_221380) regulates both translation and localisation of TgAlba2 (TGME49_218820) through interactions with the TgAlba2 3’ UTR (Gissot *et al*, 2013). Note that due to the inconsistent naming conventions for Alba domain proteins, TgAlba2 is not homologous to PfAlba2 but rather is more closely related to PfAlba3 (PF3D7_1006200) (Jagadeesh & Vembar, 2024). TgAlba1 however does belong to the same Rpp25-like grouping as PfAlba1. Together, these observations suggest that regulatory relationships between Alba family members may be conserved across apicomplexan parasites, although direct regulation of PfAlba1 expression by PfAlba2 remains to be established.

In this study, we successfully generated two endogenously tagged PfAlba2 parasite lines, enabling the characterisation of its localisation, functional importance, protein associations, and RNA targets during *P. falciparum* asexual blood stage development. We also established a customised split-TurboID as a robust, sensitive, efficient and high-confidence system for characterising the proxiome of *P. falciparum* blood stage proteins, particularly for proteins where conventional miniTurboID tagging has proven challenging. Conditional mislocalisation of PfAlba2 revealed a modest but significant growth defect following ring-stage induction, supporting its importance in normal parasite development. Complementary proximity proteomics and RIP-seq identified associations between PfAlba2, components of the translational machinery, and transcripts involved in RNA metabolism and merozoite invasion, suggesting a potential role in post-transcriptional regulation. Furthermore, the identification of PfAlba1 and PfAlba4 in the PfAlba2 proxiome, together with the association of PfAlba1 mRNA, raises the possibility of coordinated regulatory functions among Alba family members. Collectively, these findings provide new insights into the molecular functions of PfAlba2 during asexual blood stage growth and establish split-TurboID as an additional tool for investigating protein associations in malaria parasites.

## Methods

### Continuous culture of *P. falciparum* parasites

*P. falciparum* parasites were maintained as previously described (Trager & Jensen, 1976). Parasites were grown at 3% haematocrit in O+ human red blood cells (Lifeblood, Australia) in RPMI-HEPES supplemented with hypoxanthine (367 µM), gentamicin (25 µg/mL), NaHCO_3_ (0.17%), and Albumax II (0.5%) at a pH of 7.4 and stored in atmospheric conditions of 1% O_2_, 5% CO_2_, and 94% N_2_ at 37°C.

For proximity labelling assays, parasites were cultured in biotin-free RPMI media supplemented with sodium bicarbonate (1% w/v), Albumax II (0.5%), gentamicin (25 µg/mL), HEPES (1M), and hypoxanthine (367 µM) for at least two rupture cycles under standard gas and incubation conditions.

### Construction of transfection plasmids for endogenous tagging

For details of all primers used in this study, see Supplementary table 4.

The selection linked integration (SLI) method was used for endogenous tagging of target genes and plasmids were constructed as previously described (Birnbaum et al., 2017). Primers were used to insert restriction sites and PCR amplify C-terminal homology regions (HR) of PF3D7_1346300 and PF3D7_1347500 from 3D7 wildtype gDNA.

For 2xFKBP-GFP tagging, amplified inserts and the plasmid vector, pSLI-2xFKBP-GFP, were digested with NotI and AvrII (NEB). For split-TurboID tagging, pSLI-2xFKBP-GFP plasmids containing a 3’ HR for either PF3D7_1346300 or PF3D7_1347500 and the split-TurboID insert were digested with AvrII and MluI followed by ligation. All transfection plasmids were sequenced to confirm correct ligation prior to transfection.

### Transfection of *P. falciparum* parasites

Late-stage parasites were transfected as previously described (Moon *et al*, 2013). Briefly, cultures were synchronised to late-stage schizonts by 150 nM of the egress inhibitor ML10 then purified by 70% Percoll centrifugation (Rivadeneira *et al*, 1983).

For each transfection, 50-100 ug of plasmid DNA was ethanol precipitated, resolubilised in 10 μl of 1X TE buffer, added to 80 µl of cytomix (0.895% KCl, 0.0017% CaCl_2_, 0.076% EGTA, 0.102% MgCl_2_, 0.0871% K_2_HPO_4_, 0.068% KH_2_PO_4_, 0.708% HEPES) then added to concentrated schizonts. Parasite mixture was then moved to a BioRad cuvette (0.2 cm) and electroporated (0.31 kV/capacitance of 960 µF). Post-transfection, parasites were moved to 5 mL dishes for O/N incubation.

For integration lines, transfected ring stage parasites were treated with 5 nM WR99210 (Jacobus Pharmaceuticals) the following day to select for episome retention. Once stable episome retention was achieved, tag integration parasites were further treated with 400 µg/mL G418 (Thermofisher) for integration of the desired tag. All integration lines were assessed by PCR of genomic DNA to confirm successful integration.

Episome retention of the nuclear mislocalisation plasmid (1xNLS-FRB-mCherry-DHODH^NMD3^) was selected for by 0.9 µM of DSM1 (BEI resources) (Birnbaum *et al*., 2017; Branon *et al*, 2018b).

### Saponin lysis of parasite cultures

Parasite lysates harvested by 0.15% (w/v) saponin (ThermoFisher Scientific) lysis with 1X protease inhibitor (Cømplete, Roche) were incubated for 10 min on ice then centrifuged at 16,000 x g for 10 min/4°C. Lysates were then washed once in 0.075% (w/v) saponin/1X protease inhibitor followed by three washes in 1X PBS/1X protease inhibitor then frozen at - 80°C.

### SDS-PAGE and Western blot

Parasite lysates were treated with DNase (Qiagen), resuspended in reducing sample buffer (0.125 M Tris-HCl pH 7, 20% (v/v) glycerol, 4% (v/v) SDS, 10% (v/v) β-mercaptoethanol (Sigma-Aldrich), 0.002% (w/v) bromophenol blue (Sigma-Aldrich)) and separated on 4-12% Bis-Tris pre-cast gels (Invitrogen) in 1X MES running buffer (Bolt) 90 min at 110 V. Proteins were then transferred to nitrocellulose membranes (BioRad, Invitrogen) at 25 V for 7 min. Membranes were blocked for 1-hour RT in 3% skim milk/1X PBS. Primary antibodies were diluted in 0.5% skim milk/1X PBS-T 0.05% at the following concentrations and incubated O/N at 4C: 1:7000 mouse or rabbit anti-EXP2 (gifts from Paul Gilson), 1:1000 mouse anti-GFP (Sigma-Roche/Invitrogen), 1:5000 rabbit anti-aldolase (Invitrogen), 1:2000 mouse anti-FKBP (Santa Cruz), 1:2000 anti-streptavidin IR800CW (LI-COR Biosciences), 1:2000 rabbit anti-mCherry (BioVision).

Membranes were washed three times in wash buffer (1x PBS-T 0.05%) then secondary antibodies added at the following concentrations: 1:10,000 goat anti-mouse IR800CW (LI-COR Biosciences) and 1:15,000 goat anti-rabbit IR680RD (LI-COR Biosciences) and incubated at RT for 1-hour in the dark. Membranes were washed twice more in wash buffer, once in 1X PBS followed by imaging with a BioRad ChemiDoc imager.

All image adjustments and volume quantifications were undertaken using the BioRad ImageLab software (version 6.1).

### Mislocalisation assays

At ∼32 h.p.i. parasites were treated with 150 nM of ML10 for 10 hours then washed and allowed to rupture O/N. The following day when parasites were ∼10 h.p.i., they were further synchronised by 5% w/v D-sorbitol incubation for removal of any remaining late-stage parasites. Parasite cultures diluted to 1% parasitaemia/3% haematocrit were split across five 5 mL cultures. Mislocalisation was induced with 250 nM of Rapalog A/C heterodimeriser (Takara) at the following time points – 10 h.p.i., 19 h.p.i., 31 h.p.i., and 40 h.p.i. Culture media was replaced at 48-hours and parasites were incubated for an additional 24-hours.

### Assay analysis

Thin blood smears for were fixed in 100% methanol at each time point and stained with Giemsa (5% v/v) before imaging with a Zeiss Axiolab 5 microscope.

From end point smears, parasitaemia, life stage distribution and aberrant cell types were counted three times across separate days to ensure consistent counts.

### Immunofluorescence assays

Parasites were grown to >5% parasitaemia then diluted to 0.3% haematocrit and incubated on coverslips coated with 0.01% poly-L-lysine (Sigma-Aldrich) at 37°C for 30 min before washing with MilliQ H_2_O. Remaining adhered parasites were fixed with 4% paraformaldehyde (PFA) (Sigma-Aldrich) or 4% PFA/0.075% glutaraldehyde (Electron Microscopy Sciences) at 37°C for 20 min.

Fixed parasites were permeabilised (0.1% v/v Triton-X100 for 10 min) then blocked with 3% w/v bovine serum albumin (BSA)-PBS-T 0.05% for 1 hour RT. Primary antibodies were diluted in 1% BSA-PBS-T 0.05% to the following concentrations: 1:1000 rabbit anti-GFP (Genescript), 1:500 mouse anti-EXP2, 1:250 anti-streptavidin AF488 (Invitrogen), 1:250 mouse-anti-FKBP (Santa Cruz), 1:200 chicken anti-HA ab911 (Abcam) then incubated with coverslips O/N at 4°C. The following day, coverslips were washed in 1X PBS-Tween 20 0.05% three times then incubated for 1-hour in the dark RT with 1:1000 DAPI and secondary antibodies at the following concentrations: 1:500 goat-anti-rabbit AF488, 1:500 goat-anti-mouse AF594, 1:500 anti-chicken AF488, 1:200 STAR Green 488 in antibody diluent RT for 1-hour in the dark. Antibodies were post-fixed at room temperature for 10 min, washed once with MilliQ followed by dehydration in ascending concentrations of ethanol (70%, 90%, 100%) for 3 min each. Air-dried coverslips were mounted on glass slides with Prolong Gold or Prolong Diamond mounting medium and allowed to harden for 72-hours prior to imaging.

Imaging was undertaken with a Zeiss LSM980 fluorescence microscope with a Plan-Apochromat 63x/1.4 oil DIC objective lens and 405 nm, 488 nm, and 594 nm laser lines. Images were Airyscan processed.

### Live cell imaging

Parasites diluted to 0.3% haematocrit were supplemented with 300 nM DAPI and for dimerisation assays, 250 nM of Rapalog. iBidi 15 μ-slide eight-well glass-bottom slides were used for imaging. Wells were coated with 0.01% poly-L-lysine and incubated with 200 μL of sample per well for 1-2 hours before imaging. Imaging was performed using a Nikon Ti2E fluorescent microscope with a Plan Apochromat 60x NA 1.4 oil immersion objective lens with the following light sources: DAPI-5060C-NTE-ZERO, FITC-3540C-NTE-ZERO, and TxRED-4040C-NTE-ZERO.

### Image analysis

Microscopy images were analysed in Fiji/ImageJ software and adjusted for brightness, contrast, and/or colour balance. Images were cropped as appropriate and scale bars added.

### Culture and harvest of parasites for BioID experiments

Split-TurboID parasite lines were synchronised with sorbitol and grown in 125 mL volumes at 3% haematocrit until reaching 10-12% schizonts then further synchronised with 150 nM ML10 egress inhibitor. Media was removed and two 125 mL cultures were resuspended in biotin-free media, combined then split between 2x 30 mL culture dishes with 5 mL set aside as an untreated control for western blot validation. 30 mL cultures were supplemented with 150 µM of biotin and the experimental sample was additionally treated with 250 nM of Rapalog. Cultures were further incubated in a pre-warmed gas box for 30 min at 37°C under standard gas conditions.

Post-incubation ML10, excess biotin, and residual Albumax were removed by two washes in 1X PBS. Cultures then underwent saponin lysis as described above.

### Enrichment of biotinylated peptides

Samples were resuspended in lysis buffer (2% SDS, 25 mM Tris-HCl at pH 8, 150 mM NaCl, 250 U/mL benzonase nuclease (Millipore), MilliQ H_2_O, 1X phosphatase inhibitor (Roche), 1X protease inhibitor (Cømplete, Roche) and homogenised by sonication at 50% Amp in 30-second bursts. Samples were then diluted 6-fold in −20°C acetone (Sigma), incubated for 2-hours at −20°C to precipitate all proteins, then lysates clarified by 10-minute centrifugation at 4C/13,000 x g. Air-dried pellets were re-solubilised in 8M urea (Sigma) and 50 mM ammonium bicarbonate (ABC) (Sigma) then denatured for 1-hour at 38C/650 rpm. Protein concentrations were determined by BCA assay as per the kit protocol and concentrations normalised across samples. Each sample was reduced and alkylated using 10 mM Bond-breaker Tris(2-carboxyethyl)phosphine (TCEP) (Thermofisher) and 40 mM 2-chloroacetamide (CAA) (Sigma), and further diluted 8 times using 50 mM ABC to reach a final urea concentration of 1 M then incubated for 15 min at 55C/650 rpm. CaCl_2_ was then added to a 1 mM final concentration, incubated for a further 15 min at 45C/650 rpm then Trypsin/Lys-C (Promega) was added at a 50:1 w/w ratio and samples were incubated at 37°C/650 rpm for 16-hours.

For each sample, 100 µl of streptavidin magnetic beads (Genscript, Chameleon Science) were washed in 1 ml of wash buffer (2 M urea, 25 mM Tris-HCl at pH 8, 150 mM NaCl). Trypsinised samples were centrifuged at 10,000 x g for 3 min and supernatant was added to the prepared streptavidin beads and incubated for 4-hours RT with end-over-end rotation for enrichment of biotinylated peptides. Beads were magnetised to a rack for removal of supernatant and washed twice in 1 ml of wash buffer and once in mass spec grade H_2_O. Biotinylated peptides were eluted with 60°C/5 min incubations in elution buffer (80% acetonitrile (ACN) (Sigma), 0.2% trifluoroacetic acid (TFA) (Sigma), 0.1% formic acid (FA) (Thermofisher)) and magnetisation to remove the peptide-containing supernatant. The elution step was repeated five times followed by removal of ACN by vacuum centrifugation prior to desalting with C18 stage tips.

### Mass spectrometry analysis

Peptide samples were analysed with a Vanquish Neo nano LC system coupled with an Orbitrap Astral mass spectrometer with NanoSpray Flex source (Thermo Scientific) Aurora Ultimate 25×75 C18 UHPLC columns (IonOpticks) with a 20-30 min acquisition time..

### Analysis of BioID data

Raw mass spectrometry files were processed with PEAKS Studio 13 (Bioinformatics Solutions) and data refinement performed using default instrument-specific parameters. Database searching was conducted against the [*Homo Sapiens* SwissProt (June 2023), (UP000001450, release version 2016_04)] appended with a library of common laboratory contaminants and an in-house *P. falciparum* spectral library.

Default search parameters were used with the exception of Biotinylation (K, +226.0776 Da) which was included as a variable modification with allowance for a maximum of three variable modifications per peptide. A strict false discovery rate (FDR) threshold of 1.0% was applied at both peptide-spectrum match and protein levels. Relative quantification of the BioID was conducted directly at the peptide level. Peptides were filtered for those identified with a +226.0776 biotin modification. Intensity values for biotinylated peptides were then log10 transformed (missing values were assigned as ‘0’) and a *p-*value generated by students t-test. A protein level *p-*value was then obtained for each biotinylated protein from the harmonic mean of individual peptide *p-*values. Fold change of biotinylated peptides for each protein was calculated as total biotinylated peptides in +Biotin/+Rapalog samples / total biotinylated peptides in +Biotin only control samples and log2 transformed. Proteins that met the following cut-offs were considered significantly enriched: minimum biotinylated peptide log2 fold change of 2 in +Biotin/+Rapalog samples and A) appeared across two or more +Biotin/+Rapalog replicates and were absent in the +Biotin only control or B) biotinylated peptides were present in the +Biotin control but were present in at least 3 +Biotin/+Rapalog replicates.

### PfAlba2 protein immunoprecipitation and RIP-seq

Schizont-stage parasites from the PfAlba2 FKBP-GFP transgenic line were harvested in three independent biological replicate experiments for RNA immunoprecipitation sequencing (RIP-seq). Each replicate comprised approximately 10–15 × 10^6 tightly synchronised late-schizont parasites. Infected erythrocytes were lysed with 0.05% (w/v) saponin in ice-cold 1× PBS supplemented with protease inhibitor for 2–3 min on ice. Parasites were collected by centrifugation at 16,000 × g for 5 min at 4 °C and washed three times with ice-cold 1× PBS containing protease inhibitor

Washed parasite pellets were resuspended in 10 pellet volumes of extraction buffer containing 1% NP-40, 150 mM HEPES and 150 mM NaCl (pH 7.0), supplemented with protease inhibitor. Samples were sonicated on ice for 30 s at 20% amplitude using 1-s on/1-s off pulse cycles, incubated overnight at 4 °C with rotation, and clarified by centrifugation at 16,000 × g for 20 min at 4 °C.

Ten per cent of each clarified lysate was retained as the unenriched total-RNA input control. The remaining lysate was incubated with 20 μL anti-GFP magnetic beads for 2 h at 4 °C with gentle rotation. Beads were collected using a magnetic rack and washed sequentially five times with 1 mL extraction buffer, five times with 1 mL 0.5× extraction buffer and five times with 1 mL nuclease-free water. Washed bead-bound complexes were subjected to proteinase K digestion before RNA extraction.

For input samples, 100 μL of the reserved lysate was mixed with 1 mL TRIzol reagent. Immunoprecipitated RNA was extracted by directly resuspending the washed beads in 1 mL TRIzol following proteinase K digestion. Samples were mixed with 200 μL chloroform, vortexed briefly and centrifuged at 13,000 × g for 15 min at 4 °C. RNA was precipitated from the aqueous phase by addition of 500 μL ice-cold isopropanol and incubation for 10 min at room temperature, followed by centrifugation at 12,000 × g for 10 min at 4 °C. RNA pellets were washed once with 1 mL ice-cold 70% ethanol, centrifuged at 7,500 × g for 5 min at 4 °C, air-dried and resuspended in 50 μL nuclease-free water.

RNA was treated with TURBO DNase according to the manufacturer’s protocol. Briefly, 0.1 volumes of 10× TURBO DNase buffer and 1 μL TURBO DNase were added to each sample, followed by incubation at 37 °C for 20–30 min. DNase inactivation reagent was subsequently added, samples were incubated for 5 min at room temperature and centrifuged at 10,000 × g for 1.5 min, and the RNA-containing supernatant was transferred to a fresh tube. RNA concentration was determined using a Qubit fluorometer with the RNA High Sensitivity assay, and RNA integrity was assessed using an Agilent TapeStation.

### Library preparation, sequencing and read processing

Input and PfAlba2 RIP RNA libraries were prepared using the GenScript Ultra-Low Input mRNA Non-Directional Library Prep Kit and sequenced as 150-bp paired-end reads on an Illumina MiniSeq. Raw read quality was assessed using FastQC. Reads were trimmed using Trimmomatic v0.39 with a four-nucleotide sliding window, a minimum mean quality score of 20, removal of three nucleotides from both read ends and a minimum retained length of 50 nt. Trimmed reads were aligned to the Plasmodium falciparum 3D7 reference genome (PlasmoDB release 68) using Subread, and transcript-level counts were obtained using featureCounts with the corresponding PlasmoDB annotation.

### RIP-seq enrichment analysis and target identification

For enrichment analysis, featureCounts values were combined with transcript-length information and converted independently for each library to transcripts per million (TPM). Transcripts were retained when >10 raw reads were detected in at least two of three RIP libraries and independently in at least two of three input libraries. For each retained transcript, RIP and input abundance were summarised as the median TPM across the three biological replicates, and enrichment was calculated as the RIP-to-input median TPM ratio after addition of a 1-TPM pseudocount. Enrichment values were subsequently log2-transformed.

The distribution of log2(RIP/Input) enrichment values was fitted using a two-component Gaussian mixture model. The components with the lower and higher fitted means were designated the lower-enrichment and RIP-enriched statistical populations, respectively. For each transcript, the posterior probability of membership in the lower-enrichment component was treated as a local model-based false-discovery probability. Transcripts were ranked from highest to lowest enrichment, and a GMM-derived cumulative false discovery rate (FDR) was calculated as the running mean of these probabilities.

Primary PfAlba2-associated transcripts were defined by ≥2.5-fold RIP/Input enrichment and GMM-derived cumulative FDR ≤0.05. A posterior probability ≥0.95 for assignment to the RIP-enriched component, equivalent to a natural-log posterior odds score (LOD) ≥2.94, was retained as an additional confidence measure but was not used as an independent primary target criterion. A stringent high-confidence subset was defined by ≥5-fold enrichment and GMM-derived cumulative FDR ≤0.01.

### 3′UTR motif analysis

To identify putative cis-regulatory elements associated with PfAlba2 target recognition, 3′UTR sequences were extracted from the P. falciparum 3D7 reference genome using PlasmoDB release 68 annotations. PfAlba2 target 3′UTRs of at least 50 nt were retained for analysis. Non-target 3′UTRs were obtained from transcripts evaluated in the RIP-seq experiment that were not significantly enriched and were used as the discriminative background set.

XSTREME (MEME Suite v5.5) was run in discriminative RNA mode using motif widths of 6–15 nt, with the non-target sequences used to construct the background model. Motif enrichment was assessed using SEA with Fisher’s exact test, with statistical significance reported as E-values and FDR-adjusted q-values. Discovered motifs were compared with experimentally characterised RNA-binding protein motifs in the Ray2013 RNAcompete database using Tomtom (Ray *et al*, 2013a). FIMO was used to identify individual PfAlba2-associated 3′UTRs containing selected enriched motifs.

### Functional enrichment analysis

Functional enrichment analysis of the primary PfAlba2-associated transcript set was performed using STRING v12.0. *P. falciparum* 3D7 identifiers were mapped using the default STRING resolver, with duplicate mappings collapsed and unmapped identifiers excluded. The organism-wide *P. falciparum* 3D7 proteome was used as the reference background.

Gene Ontology Biological Process, Molecular Function and Cellular Component categories, together with KEGG and Reactome pathways, were assessed using the STRING enrichment framework. Multiple-testing correction was performed using the Benjamini–Hochberg procedure, with FDR <0.05 considered significant. Enrichment magnitude was represented by STRING Strength, defined as log10(observed/expected), and visualised using bubble or lollipop plots with point size representing the number of genes contributing to each enriched term.

### Protein–protein interaction network analysis

Protein–protein interaction networks were generated using STRING v12.0 with the default evidence channels and a medium-confidence interaction threshold of combined score ≥0.4. Disconnected nodes were excluded from network visualisation. Network modules were identified using the Markov Cluster Algorithm (MCL) with an inflation parameter of 3. Node size represented network degree, edge thickness represented the STRING combined interaction score, and module membership reflected the corresponding MCL cluster.

### Comparison with stage-specific proteomes

To assess stage-specific protein detection of PfAlba2-associated transcripts, the 380 primary PfAlba2 targets were compared with schizont and wild-type Day 3 gametocyte proteomic datasets (Chisholm *et al*., 2026; Marapana *et al*., 2026). Comparisons were performed at the PF3D7 gene level after removal of transcript-version suffixes.

A target was considered detected in the schizont proteome when its corresponding PF3D7 identifier was present in the schizont proteomic dataset. For the Day 3 gametocyte dataset, proteins were considered detected when classified as identified in wild-type Day 3 gametocytes. Where quantitative Day 3 measurements were available, protein abundance was represented by the mean normalised signal across the five wild-type Day 3 samples.

PfAlba2-associated transcripts were classified as detected in both proteomes, schizont only, Day 3 gametocyte only, or neither dataset. This analysis was used to determine the proportion of PfAlba2-associated transcripts with corresponding protein-level evidence at the schizont stage and the proportion of schizont-undetected targets subsequently detected during early gametocyte development.

### Enrichment analysis

Enrichment analysis was performed using ShinyGo version 0.85.1 and default parameters (0.05 FDR cut-off, minimum pathway size = 2, redundancy removed, # of pathways to show = 20). Network analysis was performed using the K-means clustering method in StringDB (STRING v12.0).

### Statistical analysis

All graphs and statistical analyses were generated using GraphPad PRISM 11 (GraphPad Software Inc.). Experimental information and statistical tests are indicated in the relevant figure legends.

## Data Availability

Proteomics and RIP-Seq data will be made available on an online repository upon acceptance of the manuscript. All other data and reagents are included in the manuscript or available through contacting the corresponding author.

## Acknowledgements

D.W.W. and D.J.C. are ARC Future Fellows. A.R.J. is an NHMRC Investigator Fellow. This work was supported by an ARC Discovery Project (DP250104773). M. E. Z. was supported by an ARC FT24 scholarship.

